# Spatially Exploring RNA Biology in Archival Formalin-Fixed Paraffin-Embedded Tissues

**DOI:** 10.1101/2024.02.06.579143

**Authors:** Zhiliang Bai, Dingyao Zhang, Yan Gao, Bo Tao, Shuozhen Bao, Archibald Enninful, Daiwei Zhang, Graham Su, Xiaolong Tian, Ningning Zhang, Yang Xiao, Yang Liu, Mark Gerstein, Mingyao Li, Yi Xing, Jun Lu, Mina L. Xu, Rong Fan

## Abstract

Spatial transcriptomics has emerged as a powerful tool for dissecting spatial cellular heterogeneity but as of today is largely limited to gene expression analysis. Yet, the life of RNA molecules is multifaceted and dynamic, requiring spatial profiling of different RNA species throughout the life cycle to delve into the intricate RNA biology in complex tissues. Human disease-relevant tissues are commonly preserved as formalin-fixed and paraffin-embedded (FFPE) blocks, representing an important resource for human tissue specimens. The capability to spatially explore RNA biology in FFPE tissues holds transformative potential for human biology research and clinical histopathology. Here, we present Patho-DBiT combining *in situ* polyadenylation and deterministic barcoding for spatial full coverage transcriptome sequencing, tailored for probing the diverse landscape of RNA species even in clinically archived FFPE samples. It permits spatial co-profiling of gene expression and RNA processing, unveiling region-specific splicing isoforms, and high-sensitivity transcriptomic mapping of clinical tumor FFPE tissues stored for five years. Furthermore, genome-wide single nucleotide RNA variants can be captured to distinguish different malignant clones from non-malignant cells in human lymphomas. Patho-DBiT also maps microRNA-mRNA regulatory networks and RNA splicing dynamics, decoding their roles in spatial tumorigenesis trajectory. High resolution Patho-DBiT at the cellular level reveals a spatial neighborhood and traces the spatiotemporal kinetics driving tumor progression. Patho-DBiT stands poised as a valuable platform to unravel rich RNA biology in FFPE tissues to study human tissue biology and aid in clinical pathology evaluation.

## Introduction

Spatial transcriptomics is revolutionizing our understanding of developmental biology, oncology, and disease pathology, mapping intricate gene expression patterns within their native tissue context^1^. This innovation is instrumental in discerning subtle nuances of transcriptional diversity, cellular pathways, and microscopic niches within complex tissues^2^, promising to refine diagnostic precision and guide the creation of targeted treatment modalities^3,4^. However, to date, the scope of the field primarily revolves around the analysis of messenger RNA (mRNA) expression. The transcriptome in eukaryotic cells is a myriad of all dynamic and diverse RNA molecules encompassing not only mature mRNAs encoding protein synthesis but also splicing variants, small RNAs, and other non-coding RNAs with regulatory functions^5,6^. Thus, spatial profiling of different RNA species throughout their life cycle is imperative for dissecting the full picture of RNA biology in complex tissues.

Formalin-fixed paraffin-embedded (FFPE) tissues are essential in clinical practice, being the backbone of human disease histopathological diagnoses^7^. Serving as the conventional method for preserving surgical pathology samples, FFPE processing maintains tissue morphology and cellular integrity at room temperature and is more economical than fresh frozen specimens due to storage, space, and personnel costs considerations. Clinical pathology researchers have accrued vast collections of FFPE blocks over time, creating a rich, yet underutilized compendium of materials that, accompanied by comprehensive clinical data, stands as a treasure trove for human biology and translational research^8^.

Nevertheless, FFPE specimens pose certain challenges. The RNA within these samples is susceptible to fragmentation during the paraffin-embedding process and may further experience heightened degradation under suboptimal storage conditions. Additionally, RNA may undergo chemical modifications, resulting in fragmentation or resistance to the enzymatic reactions required for sequencing. The loss of poly-A tails introduces another layer of complexity, restricting the utility of oligo-dT primed reverse transcription. Consequently, options for spatially profiling RNA molecules in this challenging tissue type are limited. While imaging-based platforms like MERSCOPE (Vizgen), CosMx (Nanostring), and Xenium (10x Genomics) have demonstrated spatial mapping of hundreds or thousands of genes, they are constrained by their reliance on a targeted panel for gene expression analysis with limited discovery power to profile diverse RNA species or unknown sequences. Similarly, the Visium (10x Genomics) chemistry for FFPE samples also relies on a predefined panel to target and capture RNA fragments, enabling near transcriptome-level measurement of gene expression, yet still confined to quantifying the expression of known protein-coding genes rather than base-by-based sequencing of RNA^9^.

In this evolving landscape, we present Patho-DBiT not only enabling spatial full-coverage base-by-base whole transcriptome sequencing but also meticulously crafted to address the distinctive challenges of clinically archived FFPE tissues (Figure 1A). Patho-DBiT integrates *in situ* polyadenylation^10,11^, deterministic barcoding in tissue (DBiT) using microfluidic chips^12^, and computational innovations to navigate and decode the rich RNA biology inherent in FFPE samples. The platform adeptly capitalizes on RNA fragmentation naturally occurring in FFPE specimens and appends poly(A) tails to a broad spectrum of RNA species, thereby overcoming traditional barriers associated with FFPE samples and even outperforming the assays conducted with fresh frozen tissues. By spatially barcoding intact mRNAs, fragmented mRNAs lacking poly(A) tails, various forms of large and small non-coding RNAs, splicing isoforms, and precursor RNAs carrying single nucleotide variations (SNVs), we now can gain a deeper appreciation of high-sensitivity transcriptomics, alternative splicing, genetic variation profile, microRNA-mRNA regulation, and RNA dynamics within complex tissues. Patho-DBiT represents a powerful technology for exploring spatial RNA biology in FFPE tissues, promising valuable insights into human disease development and biomarker discovery beyond gene expression.

**Figure 1.**
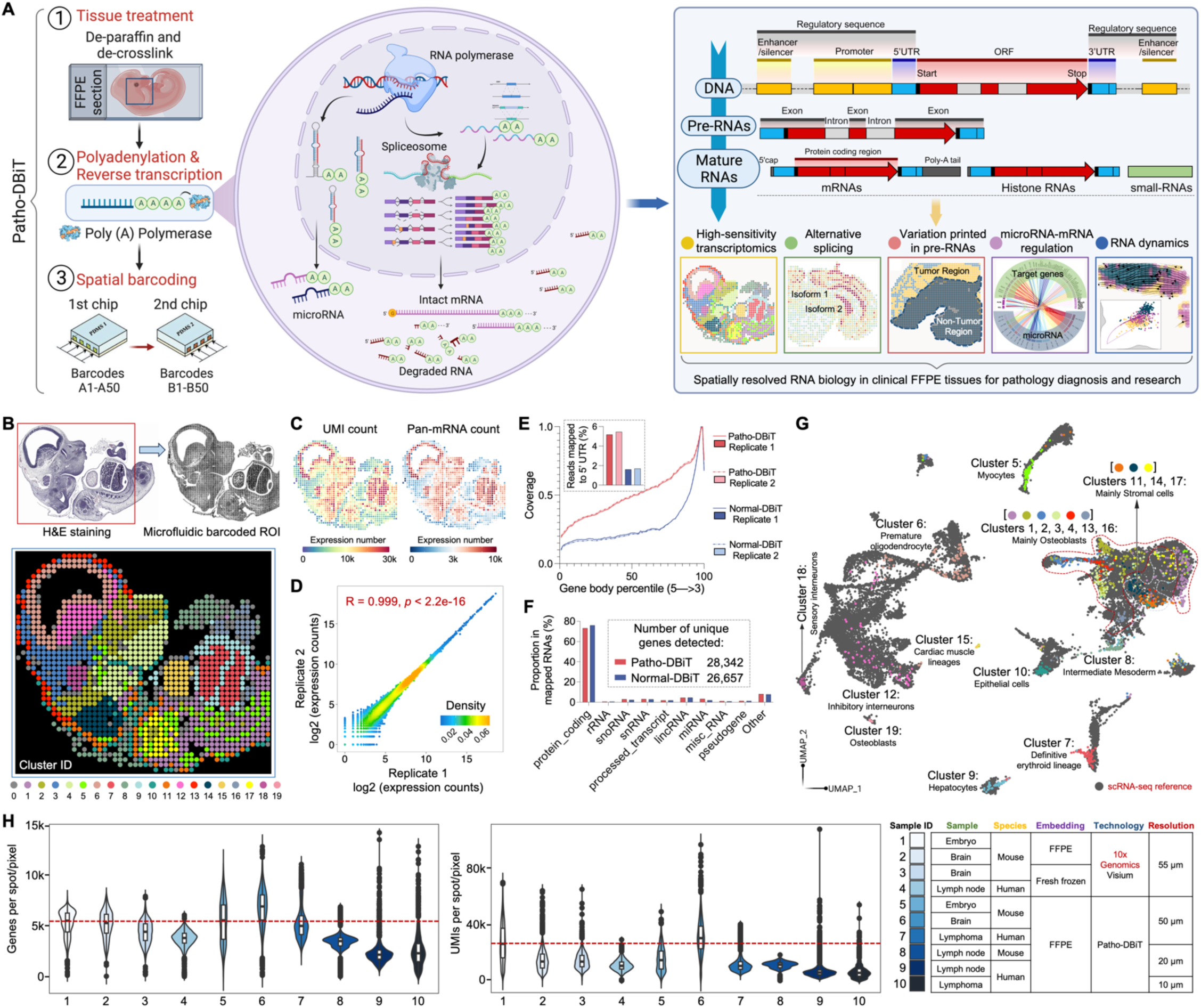
Patho-DBiT workflow, technical performance, and spatial mapping of mouse embryo. (A) Schematic workflow, molecular underpinnings, and technological spectrum of Patho-DBiT. Three major steps include (1) FFPE tissue de-paraffinization and de-crosslink, (2) Enzymatic in situ polyadenylation and reverse transcription, (3) Spatial barcoding using a pair of microfluidic devices. Patho-DBiT utilizes poly(A) polymerase to add poly(A) tails to both A-tailed intact mRNA and non-A-tailed RNAs, enabling spatial characterization of molecules across the entire transcription process. Patho-DBiT demonstrates spatial profiling of high-sensitivity transcriptome, alternative splicing, variations printed in pre-RNAs, microRNAs, and RNA dynamics. (B) Patho-DBiT’s performance and versatility on an E13 mouse embryo FFPE section. Top left: H&E staining of an adjacent section. Red square indicates the region of interest (ROI). Top right: tissue scanning post 50μm-microfluidic device barcoding. Bottom: unsupervised clustering identified 20 transcriptomic clusters, closely aligning with the H&E tissue histology. (C) Spatial pan-mRNA and UMI count maps. (D) Correlation analysis between replicates shows the high reproducibility of Patho-DBiT. Pearson correlation coefficient is indicated. (E) Read coverage along the gene body from 5’ to 3’ and the percentage of reads mapped to the 5’ UTR. Comparison involves two Patho-DBiT replicates with normal DBiT mapping without polyadenylation. (F) Comparison of the proportion of mapped RNA categories between Patho-DBiT and normal DBiT. Patho-DBiT demonstrates a similarly low level of mapped rRNA percentage compared to normal DBiT. (G) Integration of spatial RNA data with scRNA-seq mouse organogenesis data (Cao et al., *Nature* 2019). (H) Distribution of gene and UMI counts in different tissue types at varying spatial resolutions. Patho-DBiT is benchmarked against another sequencing-based spatial technology, Visium from 10x Genomics on both FFPE and fresh frozen tissues.

## Results

### Patho-DBiT design, performance, and spatial mapping of mouse embryo

The Patho-DBiT method initiates with tissue section deparaffinization and heat-induced crosslink reversal, adhering to a standardized protocol (Figure 1A). After tissue permeabilization, enzymatic *in situ* polyadenylation enables detection of the full spectrum of RNAs, followed by cDNA strand synthesis by reverse transcription. Spatial barcoding is then achieved using a microfluidic device with two PDMS chips featuring 50 parallel microchannels^12,13^. These channels sequentially deliver horizontal (A1-A50) and perpendicular (B1-B50) barcodes, creating a unique 2D barcode combination array. Post-imaging, the tissue undergoes digestion to extract barcoded cDNA to perform the downstream procedures including template switch and PCR amplification. Polyadenylation adds poly(A) tails to all RNAs, including the predominant ribosomal RNAs (rRNAs) which constitute 80-90% of cellular RNAs but provide limited information on the target transcriptome. To circumvent the loss of rare or low-abundance transcripts, cDNA fragments originating from rRNA were selectively removed from amplicons. This was achieved by employing a blend of synthetic biotinylated oligos with homology to cDNA from both cytoplasmic and mitochondrial rRNAs, resulting in the substantial reduction of these fragments prior to library sequencing (Figure S1A).

To benchmark our technology, we applied Patho-DBiT to embryonic day 13 (E13) mouse embryo FFPE section using the microfluidic device with a pixel size of 50μm. Unsupervised clustering revealed 20 transcriptomic clusters, and the spatial Uniform Manifold Approximation and Projection (UMAP) closely aligned with the histology of an adjacent section stained with hematoxylin and eosin (H&E) (Figure 1B). Cell type-specific marker genes of each individual cluster were identified, and their expression was uniquely represented in each cluster, which can be clearly separated from other clusters. Cell type-specific marker genes were identified, uniquely characterizing their expression within each individual cluster for clear distinction from other groups (Figure S1B). Notably, the distribution of these clusters exhibited conspicuous and distinctive spatial patterns, underscoring the high sensitivity and accuracy of this assay (Figure S1C). Patho-DBiT detected an average of 5,480 genes and 15,381 unique molecular identifiers (UMIs) per pixel, with the genome-wide pan-mRNA and UMI maps displaying a strong concordance with tissue morphology and density (Figure 1C). Reproducibility among replicates conducted on adjacent slices was notably high, as reflected by a Pearson correlation coefficient of 0.999. (Figures 1D and S1D). To assess the read coverage across gene bodies in our technology, we generated replicate datasets on adjacent E13 sections using normal DBiT-seq without polyadenylation. While still exhibiting a 3’ bias, Patho-DBiT showcased an approximate twofold increase in coverage across the entire gene body, and the percentage of reads mapped to the 5’ untranslated region (UTR) more than doubled (Figure 1E). This observation suggests the capture of more RNA molecules throughout the maturation cycle. Additionally, with a higher number of unique genes detected, Patho-DBiT maintained a comparably low level of reads mapped to rRNA (Figure 1F), reaffirming the efficacy of removing this undesired category.

For a more thorough assessment of data quality and to discern the cell identities within each spatial cluster, we integrated our datasets with single-cell RNA sequencing (scRNA-seq) reference data from E13.5 mouse organogenesis^14^. This integration yielded a cohesive pattern, aligning our spatial pixels with the scRNA-seq dataset in a well-conformed manner (Figures 1G and S1E). For example, cluster 1 in the spinal cord part and clusters 2, 3, and 4 located in the facial area were accurately assigned to osteoblasts, highlighting their involvement in the bone-forming process^15^. Cluster 5 cells seamlessly integrated with myocytes, mirroring their role in skeletal muscle development^16^. Clusters 6, 12, and 18 associated with the central nervous system (CNS) accurately mapped to neurons or oligodendrocytes. Cells in cluster 7, located in the liver region, uniquely integrated with the definitive erythroid lineage, marking their association with red blood cell development in this organ at the E13 developmental stage^17^. Furthermore, within the liver region, cluster 9 cells were distinctly assigned to hepatocytes, contributing to the liver’s structural integrity and function^18^. Additionally, cells in clusters 11, 14, and 17, linked to connective tissues or cartilage formation, were accurately identified as stromal cells^19^. Cells located in the heart region within cluster 15 were precisely inferred as cardiac muscle lineages. These findings reinforce Patho-DBiT’s high accuracy in cell type detection and spatial localization within the elaborate landscape of the developing mouse embryo.

To demonstrate its capability in detecting small RNA, we scrutinized the microRNA profile in the E13 mouse embryo sample. MicroRNAs, small single-stranded non-coding RNA molecules ∼22 nucleotides in length ^20^, were analyzed as a representative case. In total, Patho-DBiT detected 1063 microRNAs, peaking at 22 nucleotides in the count of mapped reads within our dataset. The average counts of UMIs and microRNAs in each pixel were 60 and 30, respectively (Figure S1F). The spatial distribution of microRNA UMI and pan-microRNA count showed a notable correlation with that of mRNA, implying coherent expression patterns between large and small RNAs within a spatial context (Figure S1G). To validate the mapping accuracy, we focused on miR-122, one of the earliest examples of a tissue-specific microRNA, constituting ∼70% of the total microRNA pool in the liver^21^. The miR-122 reads aligned to miR-122-5p, the known dominant mature strand produced from the pre-miR-122 hairpin (Figure S1H). Additionally, miR-122 exhibited a markedly higher proportional expression in the two liver region clusters, uniquely enriching its spatial distribution within this specific area (Figure S1H). We also investigated the expression landscape of the let-7 family of microRNA genes that play pivotal roles in mouse embryonic development. Patho-DBiT detected 11 out of 14 members of this family, with heterogenous expressions in different spatial clusters (Figure S1I).

We applied Patho-DBiT across diverse tissue types and spatial resolutions, yielding remarkable results. At a 50μm resolution, our approach exhibited superior performance compared to the probe-based 10x Genomics Visium for FFPE at a 55μm feature size, even surpassing its fresh frozen counterpart reliant on conventional 3’-targeted barcoding of polyadenylated RNAs. Consistently, we identified over 4,000 genes per pixel in lymphoma sections at this resolution and more than 3,000 genes in samples from mouse lymph nodes at the 20μm resolution. Notably, employing our microfluidic device with 10μm channels, we identified 2,292 genes and 6,021 UMIs from a lymphoma section at this near-cellular level. This accomplishment exceeds the capture efficiency of several state-of-the-art technologies employed on fresh frozen samples at the specified resolution, such as Stereo-seq^22^ (4.1-fold), Seq-Scope^23^ (>6-fold), and Slide-seqV2^24^ (>10-fold).

### Spatial co-profiling of gene expression, alternative splicing, and A-to-I RNA editing in the mouse brain

To further demonstrate the superior performance of Patho-DBiT as well as its capability to simultaneously map gene expression and post-transcriptional RNA processing such as alternative splicing and adenosine-to-inosine (A-to-I) RNA editing, we profiled a FFPE mouse coronal brain section. An average of 6,786 genes and 31,063 UMIs were detected per pixel, exhibiting a spatial distribution strongly correlated with the tissue histology outlined by the H&E staining of an adjacent section (Figure 2A). Clustering analysis, utilizing gene expression matrix, unveiled 15 anatomical clusters characterized by unique gene markers expressed in each subpopulation, and the spatial arrangements broadly aligned with the region annotations on a similar section from the Allen Mouse Brain Atlas (Figures 2B and S2A). Remarkably, the isocortex area was precisely deconstructed into three layers, assigning cluster 7 to layer 1-2, cluster 4 to layer 4-5, and cluster 10 to layer 6a-b^25^. The spatial expression pattern of the primary defining gene in each cluster closely mirrored the *in situ* hybridization (ISH) results for the same genes (Figure S2B), underscoring Patho-DBiT’s capacity to faithfully reflect fine tissue structures.

**Figure 2.**
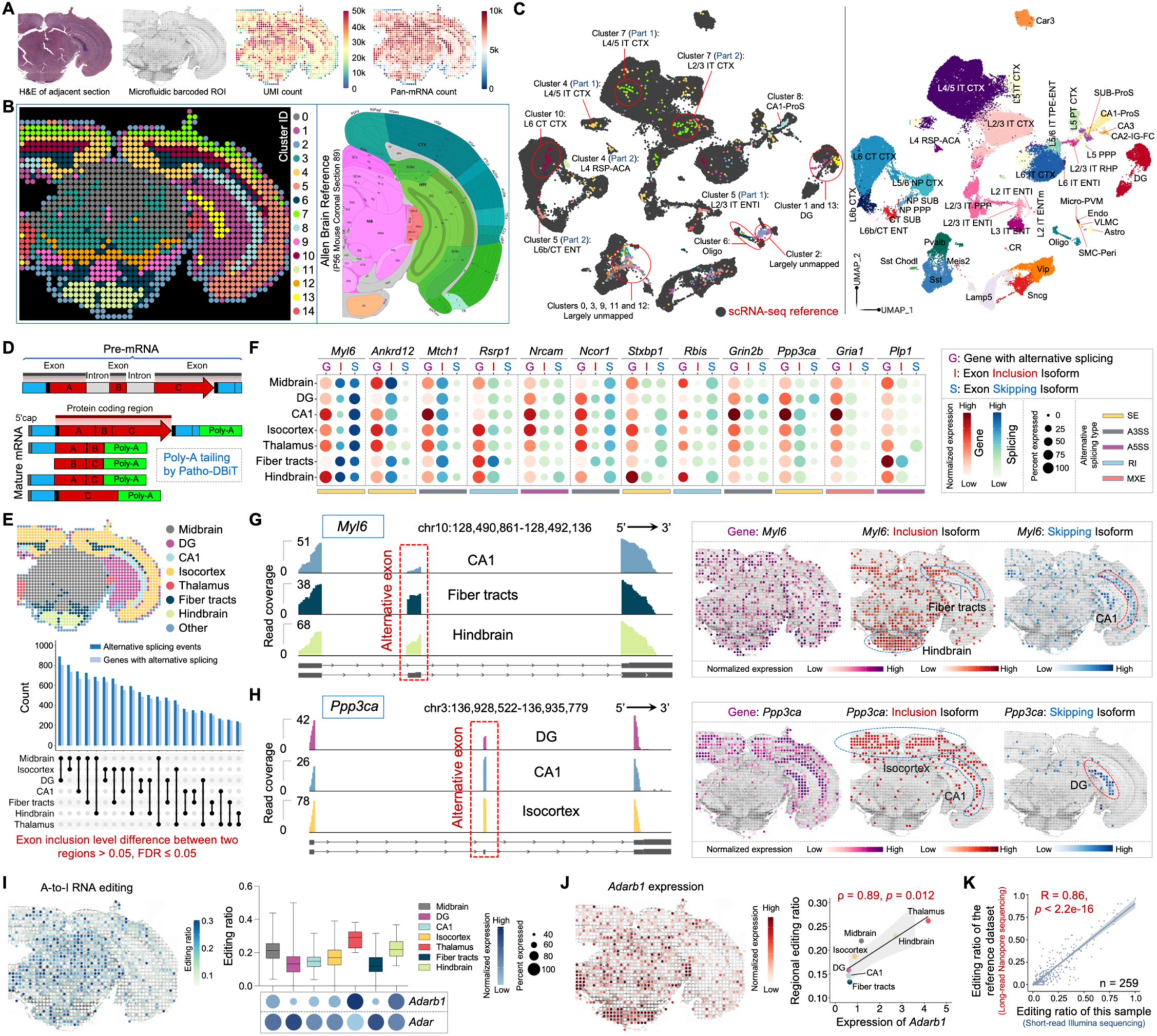
Spatial co-mapping of gene expression and RNA processing in the mouse brain. (A) Patho-DBiT profiling of an adult mouse C57BL/6 FFPE brain section. Left: H&E staining of an adjacent section. Middle: tissue scanning of the region of interest (ROI) post 50μm-microfluidic device barcoding. Right: spatial pan-mRNA and UMI count maps. (B) Unsupervised clustering identified 15 transcriptomically distinct clusters, and their distribution closely aligned with the region annotation of a corresponding coronal section from the Allen Mouse Brain Atlas (section 89, P56). (C) Integration of spatial RNA data with single-cell transcriptomics from cells in the mouse cortex and hippocampus (Yao et al., *Cell* 2021). (D) Molecular underpinnings of alternative splicing detection by Patho-DBiT. (E) Number of significant differentially spliced events and corresponding parental genes between each pair of two regions of the mouse brain. A splicing event is deemed significant if it exhibits an exon inclusion level difference > 0.05 between two regions, with a false discovery rate (FDR) of ≤ 0.05. (F) Dot plot showing the top-ranked 12 genes exhibiting significant regional differences in exon inclusion levels. Gene dot size corresponds to the percentage of pixels expressing the gene, while isoform dot size indicates the percentage of junction reads derived from the inclusion/skipping isoform over both isoforms. The color shade reflects the normalized expression level of each gene or isoform. (**G** and **H**) Junction read coverage of *Myl6* (G) and *Ppp3ca* (H) splicing event in specific brain regions. Spatial expression patterns of the gene, exon inclusion isoform, and exon skipping isoform are shown. (I) Left: spatial variations in A-to-I RNA editing in the mouse brain. Right: distribution of editing ratio across all editing sites and the expression level of ADAR-encoding genes (*Adarb1* and *Adar*) in different brain regions. Box whiskers show the minimum and maximum values. The dot size indicates the percentage of pixels expressing the gene, and the color shade represents normalized expression level. (J) Left: spatial *Adarb1* expression. Right: correlation between the *Adarb1* expression and the average reginal editing ratio across various brain regions. Spearman correlation coefficient = 0.89, *p*-value = 0.012. (K) Correlation between regional editing ratios detected by short-read Illumina sequencing-based Patho-DBiT and those detected by long-read Nanopore sequencing, as reported in the reference literature (Lebrigand et al., *Nucleic Acids Research* 2023). Analysis centered on 259 editing sites detected by both technologies, revealing a robust Pearson correlation coefficient of 0.86 (*p*-value < 2.2e-16).

Integration and co-embedding the Patho-DBiT data with the scRNA-seq atlas from cells in the mouse cortex and hippocampus validated the identity of these clusters^26^ (Figure 2C). Specifically, cells in clusters 1 and 13 integrated with the dentate gyrus (DG) type, corresponding to DG-molecular layer and DG-polymorph layer, respectively. Clusters 4, 7, and 10 consistently mapped to different layers of the isocortex, as previously described. Cluster 5 cells were also situated in the isocortex region, exhibiting a notably accurate classification as either L2/3 or L6b entorhinal area (ENT) cells. Cluster 6 cells were uniquely identified as oligodendrocytes (oligo), correlating with their distribution in the fiber tract areas. Similarly, exclusive identification was noted for cluster 8 cells, revealing their identity as hippocampal CA1 prosubiculum (CA1-Pros) cells and spatial representation. Cells in clusters 0, 3, 9, 11, and 12, located in the midbrain or hindbrain areas, remained largely unmapped due to the absence of cells from these regions in the reference scRNA-seq dataset. This provides further evidence supporting the high sensitivity and specificity of Patho-DBiT.

Alternatively spliced transcripts play a crucial role in neurogenesis and brain development, contributing to the intricate architecture of the mammalian CNS by regulating a diverse range of neuronal functions^27^. Long-read sequencing is optimal for obtaining the complete exonic structure of a transcript, but identifying splicing events from short-read RNA-seq data remains challenging despite advantages like read depth and cost efficiency^28–30^. It is due in part to the requirement of adequate read coverage for the reliable capture of the splicing-junction-spanning regions. Through the addition of a poly(A) tail during the RNA profiling process (Figure 2D), Patho-DBiT exhibited remarkably broader coverage across the gene body than other methods including poly(T) capture-based Visium spatial transcriptomics approach on a fresh frozen mouse brain section (Figure S2C). We detected a total of 3,879 distinct alternative splicing events in 2,368 genes, covering all the major event types including skipped exon (SE), retained intron (RI), alternative 3’ splice site (A3SS), alternative 5’ splice site (A5SS), and mutually exclusive exons (MXE) (Figure S2D). On average, each spatial spot yielded 105 alternative splicing events in 85 genes, with a mean coverage level of 43 UMIs per event (Figure S2E). This performance on FFPE sections is 2.2-fold higher compared to the Visium counterpart on frozen sections. In contrast, the probe-based Visium FFPE solution showed limited detection of alternative splicing events, likely attributed to its discrete capture despite a relatively broad coverage (Figures S2C and S2D).

We explored alternative splicing events with significant changes across brain regions to unravel their spatial organizational differences. Utilizing criteria set by rMATS^28^, an exon inclusion level difference > 0.05 between two regions with a false discovery rate (FDR) of ≤ 0.05 were deemed as significant. We identified a minimum of 220 differential alternative splicing events between two brain regions (specifically observed between hindbrain and thalamus), with the splicing pattern of midbrain and DG exhibiting the most pronounced differences compared to other pairs, potentially linked to their highly specialized functions within the complex brain system (Figure 2E). SE is the most abundant event type showing significant differences across regions (Figure S2F). We further delved into the top-ranked genes that are involved in brain functions and have pronounced regional isoform switching (Figure 2F). For example, *Myl6*, a gene widely involved in neuronal migration and synaptic remodeling^31,32^ with a uniform distribution of gene expression across the entire section, exhibited an enriched inclusion isoform in the fiber tracts and hindbrain, in contrast to the skipping isoform enriched in CA1 (Figure 2G). *Ppp3ca* has been identified as a leading modulator of genetic risk in Alzheimer’s disease^33–35^. Our data unveiled distinct isoform usage patterns, with the inclusion isoform prevailing in the isocortex and CA1, while the skipping isoform was exclusively expressed in the DG (Figure 2H). Patho-DBiT also identified notable examples of spatial isoform distribution, including *Nrcam* and *Stxbp1*, functional genes regulating neural development and disorders^36^ and neurotransmitter release^37^, respectively (Figures S2G and S2H).

Transcriptomic diversity expands through A-to-I RNA editing, an essential process vital for proper neuronal function^38^. With superior read coverage, Patho-DBiT spatially mapped A-to-I editing *in situ* on FFPE brain sections, unveiling a distinctive editing ratio landscape across different regions (Figure 2I). In line with prior findings^39^, conspicuous variations emerged, with thalamus exhibiting a notably elevated editing ratio (mean 27.9%), while fiber tracts displayed a lower ratio (mean 12.7%). This pattern closely corresponds to the expression levels and spatial frequencies of genes dedicated to encoding A-to-I editing enzymes, known as adenosine deaminases acting on RNA (ADARs)^40^. Particularly, the spatial distribution of *Adarb1*, known to be tissue-restricted and highly expressed in the brain^41^, closely mirrored the editing ratio, yielding a robust Spearman correlation coefficient of 0.89 (Figure 2J). Intriguingly, *Adar* showed ubiquitous expression without correlation to RNA editing (Figure S2I). To validate the accuracy of these findings, we cross-referenced the editing ratio with a published dataset generated from fresh frozen coronal brain sections using long-read nanopore sequencing^42^. In addition to the robust correlations in regional *Adarb1* expression and editing ratio between the two technologies (Figures S2J and S2K), we observed a substantial Pearson correlation score of 0.86 across 259 commonly detected editing sites showing at least 10 UMIs in both datasets (Figures 2K and S2L; Table S1). Therefore, Patho-DBiT provides a comprehensive spatial delineation of the transcriptome, alternative splicing, and A-to-I editing in the FFPE mouse brain section.

### Patho-DBiT recapitulates underlying lymphomagenesis biology in clinical archival AITL sample

We next extended the spatial profiling to clinically archived FFPE tissues. Patho-DBiT was employed to barcode a tissue section obtained from the subcutaneous nodule of a 73-year-old patient diagnosed with angioimmunoblastic T-cell lymphoma (AITL) affecting multiple lymph nodes and subcutaneous sites (Figure S3A). This block had been stored at room temperature for over five years prior to our assay. Unsupervised clustering of the gene-by-pixel matrix revealed 10 spatially organized clusters aligned with histological structures (Figure 3A). The UMAP, with an average of 5,364 genes and 11,989 UMIs per pixel, delineated distinct cell types defined by canonical markers (Figures 3B and S3B). Notably, cluster 5 exhibited a high expression of connective tissue cell-related genes (*COL1A1*, *LUM*, *FN1*), exclusively distributed within the region displaying consistent morphology observed in the H&E staining.

**Figure 3.**
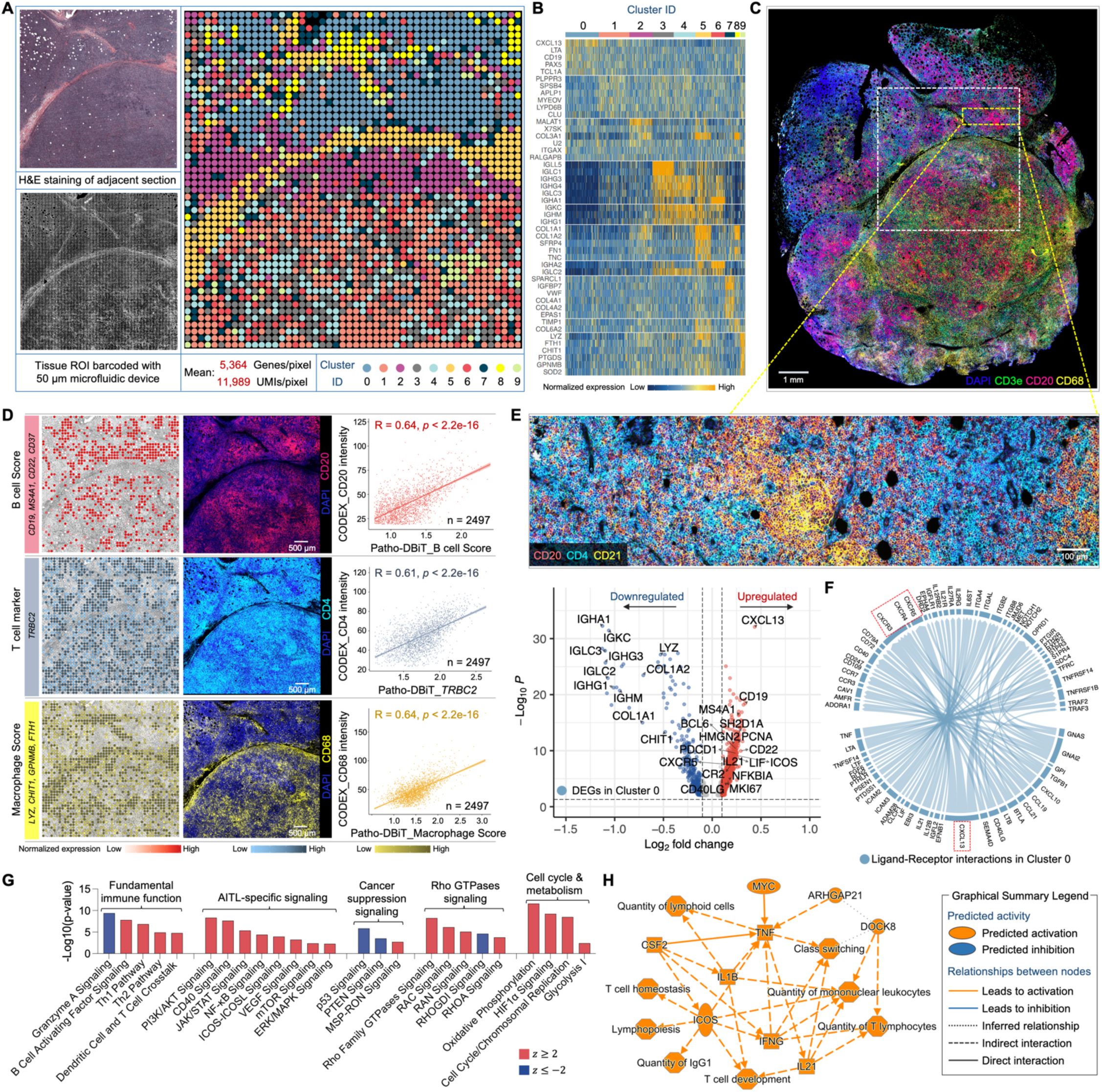
High-sensitivity spatial transcriptomics of a AITL sample stored for five years. (A) Spatial transcriptome mapping of a subcutaneous nodule section from a patient diagnosed with AITL. The FFPE block has been stored at room temperature for five years before the Patho-DBiT assay. Left top: H&E staining of an adjacent section. Left bottom: tissue scanning post 50μm-microfluidic device barcoding. Right: unsupervised clustering revealed 10 distinct clusters, aligning closely with the H&E tissue histology. (B) Heatmap showing top ranked DEGs defining each cluster. (C) Spatial phenotyping of an adjacent section using the CODEX technology (Co-Detection by Indexing). White square indicates the region of interest (ROI) in (A). (D) Spatial distributions of B cells, T cells, and macrophages revealed by Patho-DBiT, exhibiting a strong Pearson correlation with the proteomic data generated from CODEX. Genes defining each module score are listed. (E) Top: CODEX data from the yellow square indicated area in (C) showing active expression of B cell marker (CD20), T follicular helper cell (Tfh) marker (CD4), and follicular dendritic cell marker (CD21). Bottom: Volcano plot of DEGs in Cluster 0 corresponding to the indicated region. (F) Ligand-receptor interactions within Cluster 0. The distinctive communication pattern between *CXCL13* and its receptor genes (*CXCR3*, *CXCR4*, and *CXCR5*) is indicated. Edge thickness is proportional to correlation weights. (G) Corresponding canonical signaling pathways regulated by the DEGs in Cluster 0. *z* score is computed and used to reflect the predicted activation level (*z*>0, activated; *z*<0, inhibited; *z*≥2 or *z*≤−2 can be considered significant). (H) Graphical network of canonical pathways, upstream regulators, and biological functions regulated by DEGs identified in Cluster 0.

To assess the transcriptomic capture accuracy of Patho-DBiT in this long-term stored FFPE section, we conducted high-plex immunofluorescence spatial cell typing on the adjacent tissue section using the CODEX (Co-Detection by Indexing) assay^43^ (Figures 3C and S3C). The resulting cellular proteomic data faithfully depicted the histological profile of the AITL section, showcasing a population of malignant CD4+ T follicular helper (Tfh) cells intermixed with a diverse infiltrate of various immune cells^44^. Flow cytometric analysis revealed an abnormal immunophenotype of the Tfh cells: CD3+ CD4+ CD8-CD7dim+ CD5+ CD2+ CD10+ (data not shown), consistent with the extensive expression of CD4 and the near absence of CD8 and Granzyme B in the CODEX data (Figure S3D). We then traced the expression patterns of B cells (defined by *CD19*, *MS4A1* encoding CD20, *CD22*, and *CD37*), T cells (defined by the T cell receptor beta constant region gene *TRBC2*), and macrophages (defined by *LYZ*, *CHIT1*, *GPNMB*, *FTH1*). This was correlated with the surface markers of these cells—CD20 for B cells, CD4 for T cells, and CD68 for macrophages—extracted from the CODEX data. Our analysis revealed a notable correlation between the two modalities, evident in their spatial distributions (Figure 3D). The proliferation marker *MKI67* and *PDCD1*, a marker frequently expressed on malignant AITL cells, exhibited dense nodular distribution in this section, consistently identified by CODEX (Figure S3E).

Analyzing the differentially expressed genes (DEGs) within cluster 0 in comparison to the others, we observed a notable upregulation of B cell markers (*CD19, CD22, MS4A1*), markers specific to malignant Tfh cells (*CXCL13, BCL6, IL21, ICOS, CXCR5, SH2D1A* encoding SAP), and markers associated with follicular dendritic cells (*CR2* encoding CD21, *CXCL13, LIF–*IL-6 class cytokine), which are known to be abnormally proliferative in AITL^45^. The detailed profile from the high-power CODEX imaging of this region corroborated the active expression of CD20, CD4, and CD21 (Figure 3E). This intricate composition, evocative of AITL’s distinctive tumor microenvironment, warrants further investigation. We next explored cell-cell interactions by examining the expression patterns of ligands and receptors within this region. Our analysis highlighted that the most prevalent interaction pair in this network was the communication link between *CXCL13* and its receptor genes, including *CXCR3*, *CXCR4*, and *CXCR5* (Figure 3F). Notably, CXCL13 serves as a specific diagnostic marker for AITL, given its high expression in nearly all cases, and its interaction with CXCR5 is deeply implicated in tumorigenesis^44^. Patho-DBiT accurately unveiled this regulatory mechanism of AITL, complemented by the spatial distribution profile (Figure S3F).

We then delved into signaling pathways governed by DEGs within cluster 0. Alongside the activation of numerous fundamental T cell immune functions in this cluster (Figure S3G), our analysis pinpointed the upregulation of a cascade of AITL-specific signaling pathways integral to the pathogenesis process^46^, including PI3K/AKT, CD40, JAK/STAT, NF-κB, ICOS-ICOSL, VEGF, mTOR, and ERK/MAPK (Figure 3G). Conversely, general cancer suppression signals like p53 and PTEN were significantly inhibited within this cluster. Moreover, the signaling of Rho GTPases, known for their functional involvement in the initiation and progression of this specific type of T cell lymphoma^47^, showed significant activation within these cells. This activation coincided with upregulated pathways associated with cell cycle regulation and metabolism. Together, the high-sensitivity spatial transcriptomics enabled by Patho-DBiT profiling recapitulates the underlying biology of lymphomagenesis in a 5-year archived AITL tissue. This comprehensive understanding is summarized through a graphical network encompassing canonical pathways, upstream regulators, and biological functions (Figure 3H).

### Genome-scale spatial sequence variant profiling for tumor discrimination

To further assess the potential utility of Patho-DBiT in aiding pathological evaluation, we conducted spatial profiling on a biopsy from a patient diagnosed with extranodal marginal zone lymphoma of mucosa-associated lymphoid tissue (MALT), a low-grade non-Hodgkin B-cell lymphoma^48^. The section, derived from a FFPE block stored for three years at room temperature, underwent spatial barcoding using a microfluidic device with a pixel size of 50μm. Biopsy from the gastric antrum of the stomach showed a dense nodular infiltrate of lymphocytes primarily in the mucosa, as detected by endoscopy (Figure S4A). The infiltrate is composed of small to intermediate sized lymphocytes with monomorphic ovoid nuclei, condensed chromatin, and a small amount of cytoplasm. There are numerous small mature plasma cells in the lamina propria. The epithelium is predominantly intact and shows no dysplasia, and the lymphoma does not extend to the base of biopsy.

Unsupervised clustering and UMAP visualization revealed 9 clusters that spatially mirrored the histological structures (Figures 4A and S4B). Within these clusters, distinct cell types such as B cells, macrophages, plasma cells, and mucus-secreting cells were delineated based on canonical marker gene expression (Figure 4B). These cell types were uniquely distributed in Cluster E1, E4, E5, and E6, respectively. We noticed a faint expression of the Plasma Cell Score and Macrophage Cell Score in the designated Region P and Region M in Figure 4B. To validate that this signal reflects actual cellular presence rather than background noise, we conducted immunofluorescence (IF) assays targeting the immunophenotypic markers CD138 for plasma cells and CD68 for macrophages on adjacent sections (Figure 4C). The results confirmed the cell identity, providing further support for Patho-DBiT’s capability to capture rare cell types in specific regions.

**Figure 4.**
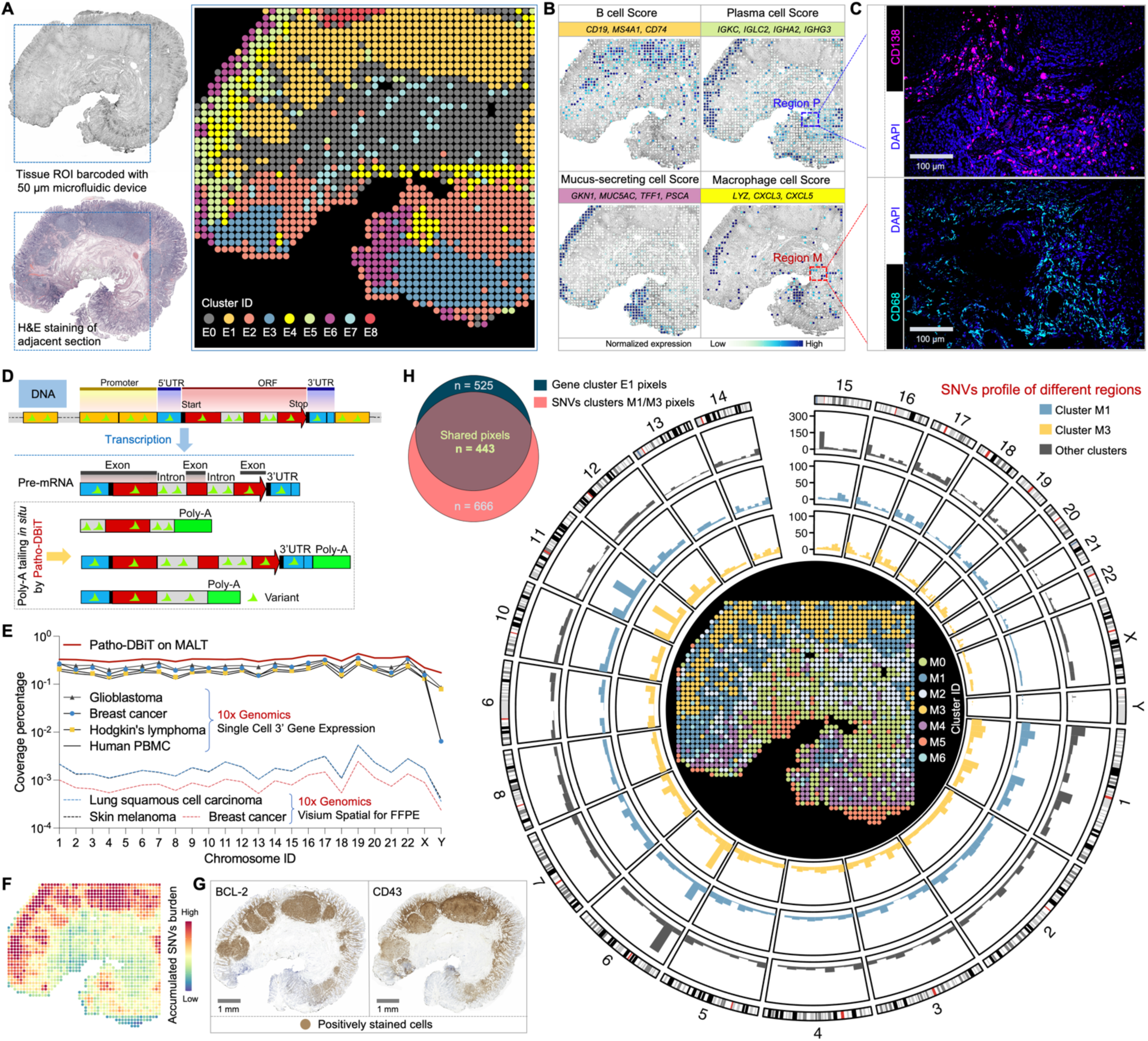
Patho-DBiT enables spatial variant profiling for tumor discrimination. (A) Spatial transcriptome mapping of a gastric antrum biopsy section from a patient diagnosed with extranodal marginal zone lymphoma of mucosa-associated lymphoid tissue (MALT). The FFPE block was stored at room temperature for three years. Left top: tissue scanning with region of interest (ROI) indicated with blue square. Left bottom: H&E staining of an adjacent section. Right: unsupervised clustering revealed 9 distinct clusters, aligning closely with the H&E tissue histology. (B) Spatial identification of representative cell types through curated expression of canonical genes. Genes defining each module score are listed. (C) Patho-DBiT’s ability to capture rare cell types in specific regions was cross-validated through immunofluorescence (IF). The IF staining of plasma cell marker (CD138) and macrophage marker (CD68) in the selected Region P and Region M in (B) was shown. (D) Molecular underpinnings of detecting variations printed in pre-mRNA transcripts by Patho-DBiT. (E) Comparison of genomic location coverage bandwidth between Patho-DBiT and other technologies. (F) Spatial expression map of accumulated single nucleotide variants (SNVs) burden. (G) Immunohistochemistry (IHC) staining of canonical markers commonly used for detecting MALT tumor cells (BCL-2 and CD43) on adjacent sections. (H) Unsupervised clustering of the spatial SNV matrix. Left: Veen plot showing the pixel overlap between gene cluster E1 and SNV clusters M1 and M3. Right: genome-wide distribution of somatic variations in clusters M1 and M3 using pixels from the other clusters as controls. Only high-confidence variant loci were preserved for downstream analysis and visualization.

Sequence variations were often found in RNA transcripts, such as those reflecting the underlying genetic mutations and those as the consequence of RNA editing, a process frequently deregulated in cancer^49,50^. By *in situ* poly(A) tailing of the entire RNA spectrum (Figure 4D), we hypothesized that Patho-DBiT could effectively capture sequence variations printed in pre-mRNA transcripts. Higher genomic coverage will enable more reliable detection of sequence variants. We therefore first conducted a comparative analysis of the genomic location coverage bandwidth between Patho-DBiT and 10x Genomics single-cell 3’ gene expression datasets. Here, the coverage percentage for each chromosome is calculated by dividing the cumulative length of sequenced regions in annotated GENCODE^51^ transcripts by the total transcript length on a chromosome. Our spatial data from the FFPE MALT section showed higher coverage capability than scRNA-seq datasets from both human cancer samples and healthy donor peripheral blood mononuclear cells (PBMC), allowing Patho-DBiT to capture genome-wide sequence variations (Figure 4E). This performance is over 176-fold higher than that observed in Visium spatial FFPE datasets from various cancer samples.

To delineate the RNA variant profile of this MALT section, we implemented a variant calling pipeline to identify all potential SNVs. Each SNV site in a spatial pixel was cataloged and classified based on the reference and variant reads, overall yielding a matrix of detected SNVs (see Methods). The spatial heatmap of accumulated SNVs highlighted a notably higher SNV burden in the B cell region compared to other areas (Figure 4F). The tumor signature in these B cells was validated through immunohistochemistry (IHC) staining of canonical markers commonly used for detecting MALT tumor cells, namely B-cell lymphoma 2 (BCL-2) and CD43^52^, on adjacent sections (Figure 4G). The expression of these markers exhibited a strong correlation with the spatial SNV burden profile.

We explored the potential of leveraging RNA-encoded variant information for unsupervised tumor discrimination. Spatial clustering of the expressed SNV matrix alone revealed 7 subpopulations (Figure 4H, center). Cluster M1 and M3 exhibited notable overlap with the tumor region; out of 525 pixels from the tumor B cell-enriched E1 cluster, 443 pixels were assigned to M1 or M3. The UMAP visualization showcased the distinct variation profile of these two clusters compared to the other five (Figure S4C). Additionally, principal component analysis (PCA) demonstrated that pixel points from M1 and M3 clusters could be differentiated from the rest using the top 5 identified principal components, particularly PC1 (Figure S4D). To determine the variant profiles across different SNV clusters, we calculated the SNV frequency within sliding genomic regions of 10,000 bp, generating their genomic distribution (Figure S4E). The counts of genomic regions with SNV frequency >0.01% were significantly higher in M1 and M3, confirming their elevated variant levels (Figure S4F). To depict the genome-wide distribution of SNVs, somatic variation calling was conducted within the M1 and M3 using pixels from the other five clusters as controls, revealing a chromatin and region-specific pattern of identified SNVs within these two tumor clusters (Figure 4H). Collectively, these findings suggest that Patho-DBiT possesses the capability to autonomously distinguish tumor from non-tumor regions and potential tumor subclones based on the RNA sequence variation profile.

### Spatial regulatory network of microRNA-mRNA interactions in tumorigenesis

We next sought to assess Patho-DBiT’s capacity for co-mapping large and small RNAs in clinical samples, with a specific focus on microRNAs which play diverse roles in various pathologies including cancer^53^. Out of 2300 true human mature microRNAs^54^, Patho-DBiT detected 1808 in the MALT section, with the count of mapped reads accurately peaking at 22 nucleotides in our dataset (Figure 5A). Assessing the UMI count per pixel for all identified microRNAs, 54% had fewer than 10 UMIs, 35% had 10-100 UMIs, and the remaining 11% had more than 100 UMIs.

**Figure 5.**
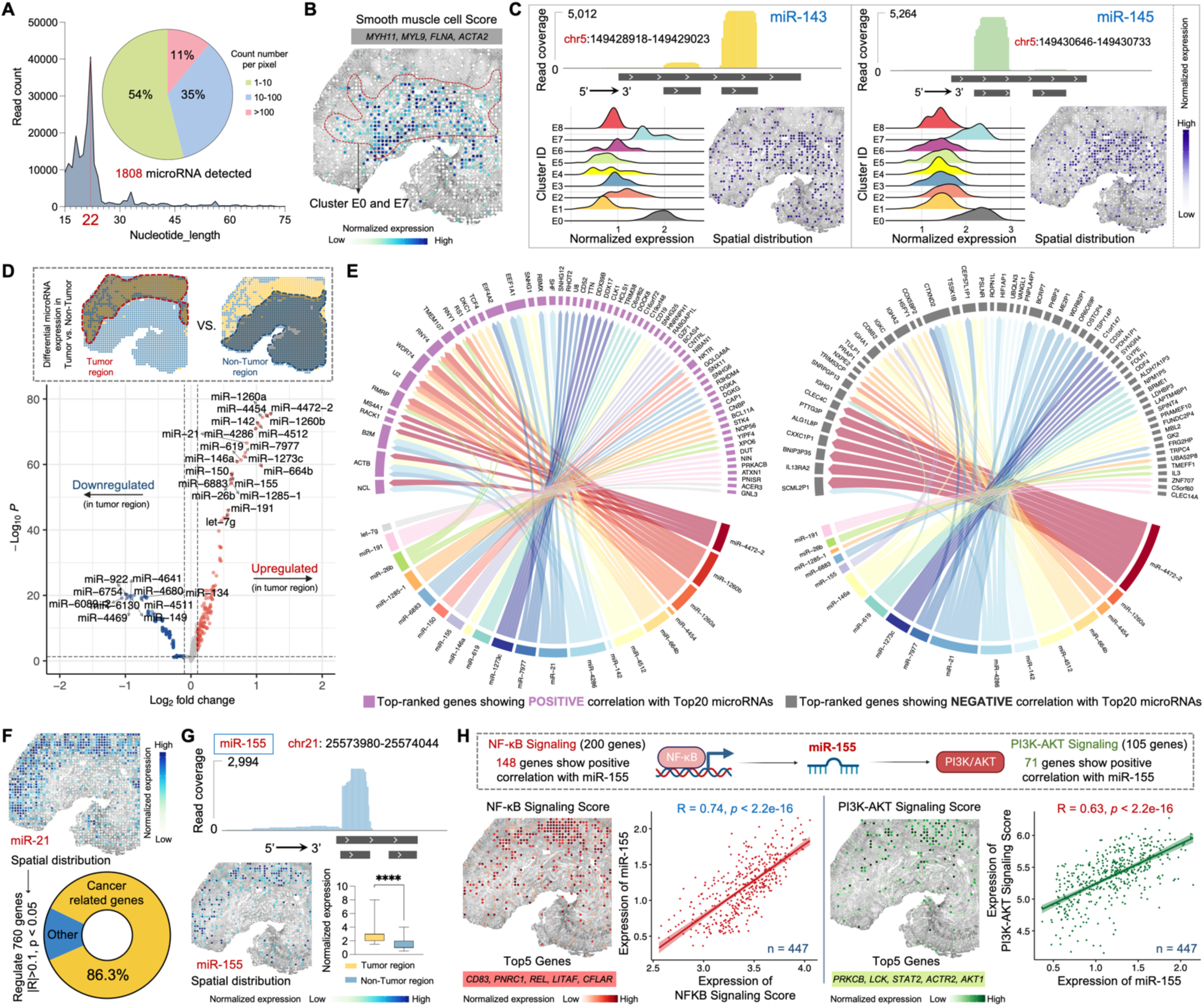
Spatial microRNA-mRNA regulatory network in the MALT section. (A) MicroRNAs detected by Patho-DBiT in the MALT section, with the count of mapped reads peaking at 22 nucleotides. The pie chart illustrates the percentage distribution of the detected count number per spatial pixel. (B) Spatial distribution of the Smooth muscle cell Score. Genes defining this module score are listed. (C) Spatial mapping of smooth muscle cell specific miR-143 and miR-145. The read coverage mapped to the reference genome location, expression proportion in each identified cluster, and spatial distribution are shown. (D) Volcano plot showing differentially expressed microRNAs between the tumor and non-tumor regions. (E) Regulatory network between the top 20 upregulated microRNAs and the gene expression in the tumor region. Genes with the highest rankings, demonstrating positive or negative correlations with the microRNAs, were separately illustrated. Edge thickness is proportional to correlation weights. (F) Spatial expression map of the oncomiR miR-21. This microRNA significantly regulates 760 genes (Pearson R > 0.1 or < -0.1, *p*-value < 0.05). Cancer-related genes are defined based on the IPA data base. (G) Spatial expression map of the B-cell lymphoma enriched miR-155. Top: read coverage mapped to the reference genome location. Bottom left: spatial distribution. Bottom right: expression comparison between tumor and non-tumor regions. Box whiskers show the minimum and maximum values. Significance level was calculated with two-tailed Mann-Whitney test, **** P < 0.0001. (H) Spatial interactions involving mir-155 and its upstream and downstream signaling pathways. Top 5 genes defining each module score are listed. The Pearson correlation between mir-155 expression and both signaling pathways was calculated across 447 spatial pixels within the tumor region.

We assessed detection accuracy by examining tissue-specific microRNAs. Based on both tissue morphology and the enriched expression of marker genes *MYH11*, *MYL9*, *FLNA*, and *ACTA*2, cells within clusters E0 and E7 were classified as smooth muscle cells^55^ (Figures 5B and S4A). Two smooth muscle cell-specific microRNAs, miR-143 and miR-145^56^, known for their involvement in proliferation, differentiation, and plasticity, exhibited markedly elevated expression in clusters E0 and E7, with over 5000 reads precisely mapped to the reference genome location, reflecting the dominant mature strands (miR-143-3p and miR-145-5p) from their respective pre-microRNAs (Figure 5C). Their spatial distributions were prominently evident in the smooth muscle cell region. Several members of the miR-30 family exert regulatory roles across different stages of mature B-cell differentiation^57^. Patho-DBiT successfully detected three of these members, namely miR-30b, miR-30d, and miR-30e, showcasing elevated expression particularly in the B cell cluster E1 or the plasma cell cluster E5 (Figure S5A). MiR-142 is necessary for the normal development of marginal zone B cells^58^, while both miR-146a and miR-150 are upregulated in marginal zone lymphomas^59^. Consistently, a notably high expression pattern of these three microRNAs was observed in the tumor B cell region of this MALT section (Figure S5B).

To further elucidate the regulatory role of microRNAs in tumorigenesis, we conducted a differential microRNA expression analysis between the tumor and non-tumor regions. Among the upregulated microRNAs in the tumor region is miR-21, a well-characterized cancer-promoting ‘oncomiR’^60^, along with abovementioned lymphoma-enriched microRNAs such as miR-142, miR-146a, miR-150, and miR-155 (Figure 5D). In contrast, miR-134 and miR-149, two microRNAs known to suppress the proliferation and metastasis of multiple cancer cells^61,62^, were significantly downregulated in the tumor region. Regulatory network analysis of microRNA-mRNA interactions in the tumor region revealed positive correlations between the top 20 upregulated microRNAs and multiple genes implicated in lymphomagenesis (Figure 5E), including *NCL* encoding a BCL-2 mRNA binding protein^63^, *ACTB* and *B2M*, which are frequently mutated in aggressive B-cell lymphomas^64^, and *EEF1A1*, potentially contributing to tumor initiation and progression^65^. Conversely, a broad array of genes exhibited negative correlations with these microRNAs, especially miR-21 and miR-4472-2, the latter also being identified for its role in fostering tumor proliferation and aggressiveness^66^. Likewise, interaction analysis was conducted for the top 10 downregulated microRNAs in the tumor region, elucidating how these regulations influence the transcriptomic signatures of tumor B cells (Figure S5C). For instance, the negative correlation observed between miR-134 and *CHD8*, *FTL*, *WNK1*, *TACC1*, *CDKL2*, and *TCF4* implies that these genes may play a role in promoting tumorigenesis in the MALT patient.

We performed detailed regulatory analysis by focusing on representative microRNAs, with miR-21 as the initial example. This microRNA exhibited enriched spatial expression in the tumor region and was inferred to significantly regulate 760 genes in this sample, with 86.3% of them identified as cancer-related genes according to the Ingenuity Pathway Analysis (IPA) database^67^ (Figure 5F). As an ‘oncomiR’ known for its pivotal role in the initiation and development of various B-cell malignancies^68^, miR-155 exhibited significantly higher expression in the tumor region, accompanied by precise genome location mapping and spatial distribution (Figure 5G). This microRNA, being a transactivational target of NF-κB^69^, contributes to the promotion of the PI3K-AKT signaling pathway in B-cell lymphoma^70^. Out of 200 genes in the Gene Set Enrichment Analysis (GSEA)-defined NF-κB signaling activation, 154 exhibited a positive correlation with miR-155 (Figures 5H and S5D). Similarly, a positive correlation was identified between miR-155 and genes linked to the activation of PI3K-AKT signaling, with 89 out of 105 genes showing this relationship (Figures 5H and S5E). The enriched spatial expression of both signaling pathways was specifically mapped to tumor B-cell region. The correlation between miR-155 expression and both signaling pathways was calculated across 447 spatial pixels within the tumor region, resulting in Pearson correlation coefficients of 0.74 and 0.63, respectively (p-value < 2.2e-16). Taken together, Patho-DBiT enables spatially resolved co-profiling of large and small RNAs, facilitating the analysis of a microRNA-mRNA regulatory network in the clinical biopsy.

### Spatial RNA splicing dynamics reveals the trajectory of tumor cell development

In all the aforementioned samples, Patho-DBiT primarily includes reads mapped to exonic regions derived from mature spliced transcripts. While these exonic reads yield an average of 4,131 genes and 15,726 UMIs per pixel in the MALT lymphoma section, we detected a substantial number of intronic molecules in this sample, corresponding to a mean pixel count of 7,509 genes and 22,583 UMIs (Figure 6A). This observation may be attributed to the poly(A) addition and subsequent capture in the intron regions. By aggregating both the exonic and intronic expression matrices while preserving their individual identities, we identified 14 clusters through unsupervised clustering analysis (Figures 6B and S6A), suggesting that this combined profile could enhance the refinement of intrinsic biological heterogeneities of this lymphoma sample. Particularly, while utilizing solely exonic reads led to the definition of only one B-cell cluster (Figure 4A), the merged matrix identified three clusters (C3, C4, and C6) within this tumor region using identical clustering parameters (Figure 6B). Their B-cell signature was authenticated by an enriched expression of the B-cell Score characterized by *CD19*, *MS4A1*, and *CD74*. The three clusters exhibited no prominent variations in cell cycle stages, as evidenced by their uniformly low levels of S and G2/M activities, further confirmed through sparse IHC staining of Ki67 (Figure 6C). Thus, their inherent distinctions and connections warrant deeper investigation.

**Figure 6.**
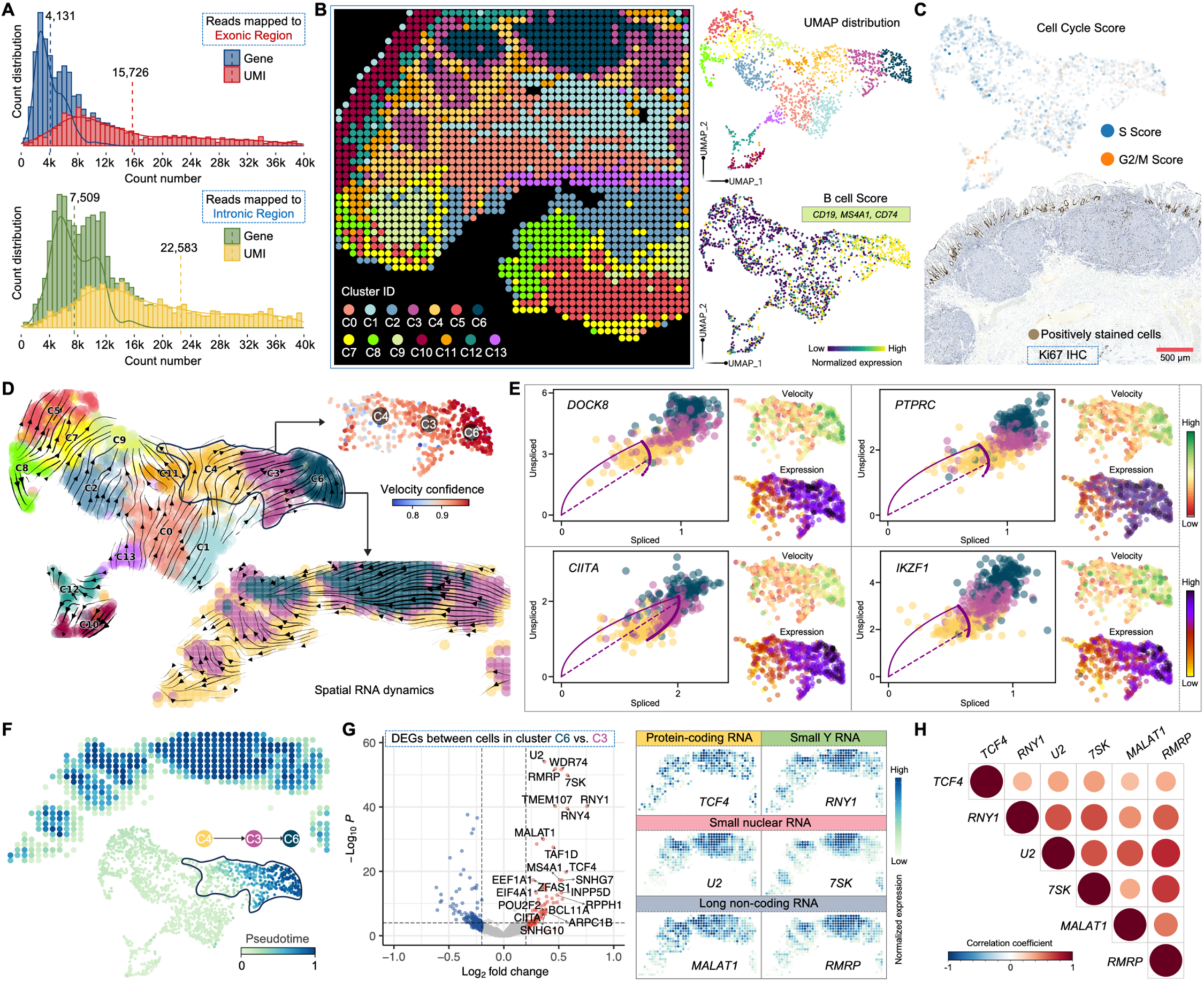
Tumor differentiation trajectory revealed by spatial RNA splicing dynamics. (A) Distribution of detected gene/UMI counts per spatial pixel from reads mapped to exonic or intronic region. The dashed lines indicate average level of gene or UMI count in the MALT section. (B) Unsupervised clustering of the combined exonic and intronic expression matrix. The analysis identified 14 clusters, showcasing UMAP visualization and featured expression of the B cell Score in clusters C3, C4, and C6. Genes defining this module score are listed. (C) Top: cell cycle score colored by the S or G2/M stage. Bottom: IHC staining for Ki67 in the tumor region of an adjacent section. (D) Velocities derived from the dynamical RNA splicing activities are visualized as streamlines in a UMAP-based embedding. The coherence of the velocity vector field provides a measure of confidence, and the spatial velocity pattern within the tumor B cells region is highlighted. (E) Phase portraits showing the ratio of unspliced and spliced RNA for top-ranked genes driving the dynamic flow from cluster C4 to C6, along with their expression and velocity level within the three tumor clusters. The dashed purple line corresponds to the estimated splicing steady state. Positive velocity signifies up-regulation of a gene, observed when cells exhibit a higher abundance of unspliced mRNA for that gene than expected in steady state. Conversely, negative velocity indicates down-regulation of the gene. (F) Spatial pseudotime of underlying cellular processes based on the transcriptional dynamics. A discernible change is evident exclusively within the three tumor clusters, where a higher pseudotime number denotes a later differentiation stage. (G) Volcano plot showing DEGs between cluster C6 and C3. Signature large and small RNAs associated with increased dynamic activities are spatially visualized. (H) Correlation matrices of the signature RNAs evaluated in G. Only significant correlations (*p*-value < 0.05) are represented as dots. Pearson’s correlation coefficients from comparisons of RNA expression across pixels in the tumor region are visualized by color intensity.

RNA velocity enables the reconstruction of dynamic changes in gene expression by leveraging the ratio of newly transcribed, unspliced pre-mRNAs to mature, spliced mRNAs, with the former identifiable through the presence of introns^6,71^. Typically, bulk or single-cell RNA-seq datasets comprise only 15–25% of reads as unspliced intronic sequences, primarily originating from secondary priming positions within intronic regions^6^. Notably, Patho-DBiT generated a higher proportion of intronic reads compared to the exonic part, enhancing the RNA velocity analysis. Employing scVelo^71^, a method that characterize the full transcriptional dynamics of splicing kinetics using a likelihood-based dynamical model, we delineated transient cellular states of all the identified clusters based on the combined matrix. Within the spatially organized tumor B-cell region, we observed a developmental trajectory originating from C4 extending towards C6 (Figure 6D). Across the majority of pixels, with high velocity confidence exceeding 0.9, we constructed a spatial RNA splicing dynamics map for MALT lymphoma tumor cells, potentially unveiling their differentiation stages as defined by splicing rates. Furthermore, we identified prominent genes that drove the primary processes of this dynamic behavior, with the top-ranked genes clearly displaying higher splicing activities in C6 compared to C3 and C4 (Figures 6E and S6B). Among these pivotal driving genes, *DOCK8* and *PTPRC* (also known as *CD45*) regulates BCR signaling and activation of memory B cells^72^, *CIITA*, *SLC38A1*, *SYK*, and *FCRL5* have been identified as prognostic factors for hematological malignancies^73–77^, *IKZF1* serves as a central regulator of lymphopoiesis^78^, and *ARHGAP44*, a gene encoding Rho GTPase activating protein 44, plays an extensive role in lymphoma initiation and progression^47^. All of them exhibited notably higher expression and velocity in cluster C6, indicating a greater level of upregulation and dynamic change compared to the relatively steady state observed in cells from C3 and C4.

Next, we inferred a universal pseudotime shared among genes that represents the cell’s internal clock. While all the clusters outside the tumor region displayed a static cell fate, changes began to manifest from C4 and progressively intensified toward C6 (Figure 6F). This spatial directional transition, coupled with the velocity flow from C4 to C6, further implies that tumor cells in C6 pixels were produced at a later stage. Putative driver genes contributing to this pseudotime trajectory were identified and ordered by their likelihoods across these clusters, among which the top-ranked gene, *BCL2*, is a key antiapoptotic regulator critical for lymphoma pathogenesis^79^ (Figure S6C). We further performed DEG analysis between cells in cluster C6 and C3, revealing a regulatory profile consisting of both large and small RNAs (Figure 6G). Among the significantly upregulated molecules in C6 compared to C3, *TCF4* primarily functions as a transcriptional activator in B-cell development^80^ and contributes to lymphoma pathogenesis^81^, *RNY1* is affiliated with the Y_RNA class and mainly involved in DNA replication and RNA stability^82^, *U2* spliceosomal small nuclear RNA (snRNA) is an essential component of the major spliceosomal machinery^83^, *7SK* is another snRNA controlling the activity of a major transcription elongation factor P-TEFb^84^, and *MALAT1* and *RMRP* are long non-coding RNAs that promote the development of various lymphomas^85,86^. All the small RNAs discussed here were accurately and substantially mapped by Patho-DBiT (Figure S6D). The heightened spatial expression of these molecules, along with their significant internal positive correlations (Figure 6H), further substantiates the increased dynamic activities within the C6 subpopulation. Together, with a superior capture efficiency for intronic reads, Patho-DBiT spatially mapped RNA splicing dynamics associated with the trajectory of malignant B cell development.

### Mapping spatial evolution of tumor progression at the cellular level

We extended our analysis by spatially mapping gastric fundus nodule biopsy sections using 10μm pixe size microfluidic devices at a cellular-level resolution. Collected from the same patient depicted in Figure 4(A) at the same time, this fundus biopsy showed diffuse large B-cell lymphoma (DLBCL), likely arising from the indolent MALT lymphoma precursor. The biopsy revealed a diffuse sheet of atypical large lymphocytes, extending from superficial glandular structures to the deep margin of the biopsy (Figure S7A). Predominantly large and pleomorphic, the tumor cells exhibited irregular nuclear contours, dispersed chromatin, and a moderate amount of cytoplasm, with frequent mitoses. Focal tumor infiltration of epithelium and background eosinophilia were observed, while the surface epithelium showed no significant abnormalities.

Tissue sections from two distinct regions were selected for spatial barcoding, detecting an exonic average pixel count of 2,292 genes and 6,021 UMIs in Region 1, and 1,507 genes and 3,466 UMIs in Region 2 (Figures 7A and S7B). Unsupervised clustering of Region 1 identified two clusters with similar phenotypes of B cells, distinguished by varying dynamic levels as indicated by the differential small RNA expression of *7SK*, *RNY1*, and *RNY3* in cluster 2^82,84^ (Figure S7C). Their tumor signature was verified by IHC staining for BCL-2 and CD43 (Figure S7D). In Region 2, a more intricate spatial organization of diverse cell types, including B cells, macrophages, and mucus-secreting cells, was identified (Figure 7B). Notably, Patho-DBiT resolved intrinsic heterogeneities among tumor B cells. While in both cluster 2 and 5, cells showed enriched expression of B-cell markers, cluster 2 displayed enhanced chemokine gene activity (Figure 7C). These genes engaged in extensive communication with cells in cluster 5 through ligand-receptor pairs (Figure S7E), potentially linked to their significant upregulation of Rho GTPases related signaling pathways^87^. Clusters 4, 7, and 8 were characterized as gastric mucus-secreting cells, exhibiting high expression of signature genes including *MUC5CA*, *TFF1*, and *PSCA*^88^. Patho-DBiT unraveled their molecular heterogeneities: cluster 7 showed elevated *PIGR* expression actively participating in the transcytosis of soluble polymeric isoforms of immunoglobulins^89^, cluster 8 displayed higher *GNK1* expression related to gastric mucosal inflammation^90^, and cluster 4 had reduced *MUC1* expression compared to the other two. These subpopulations, along with cluster 6 enriched with plasma cells, form a unique transcriptomic neighborhood revealed by our cellular-level spatial mapping, closely aligning with the tissue morphology defined by H&E staining of an adjacent section (Figure 7D).

**Figure 7.**
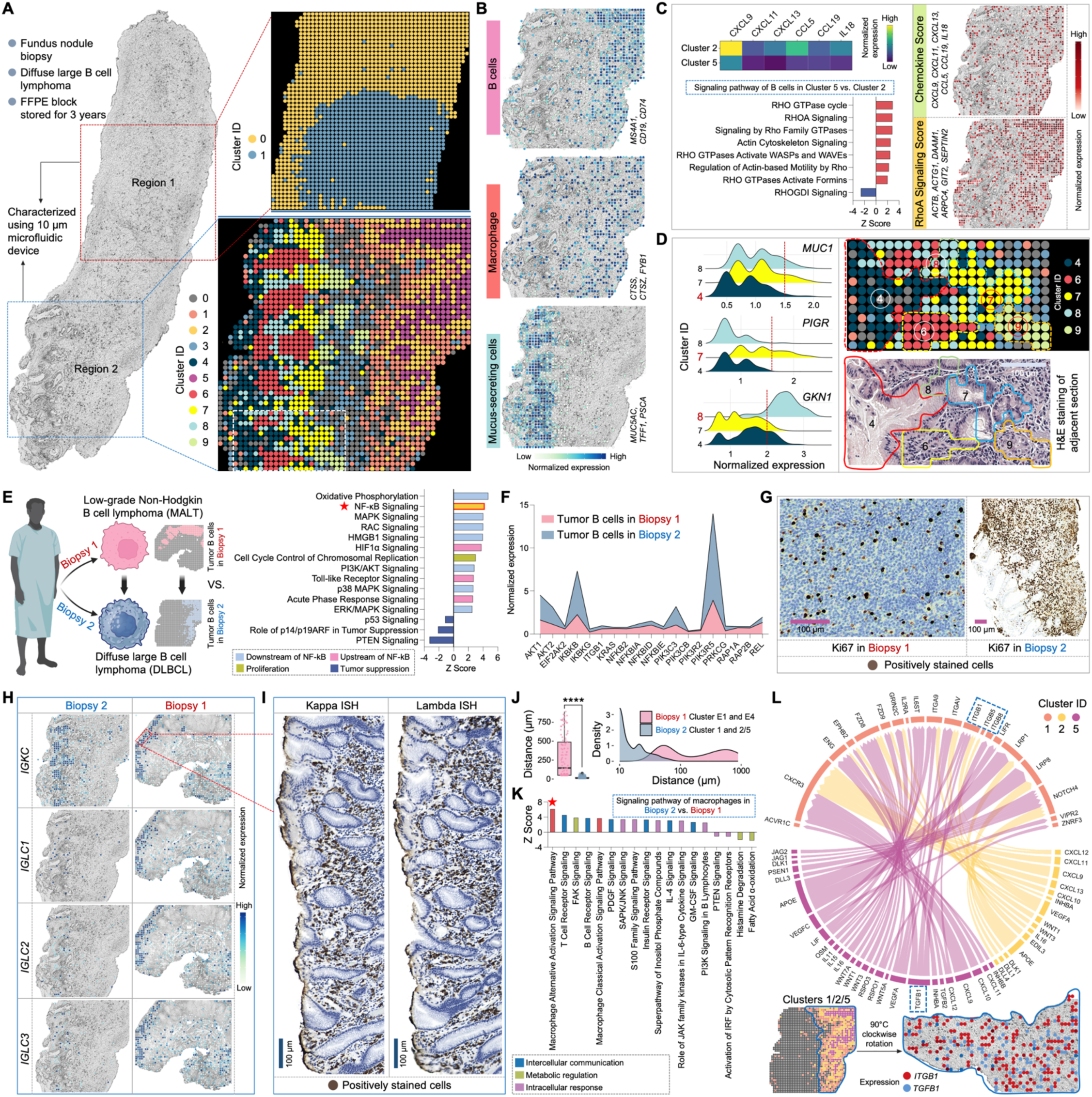
Cellular level spatial mapping of a DLBCL section elucidates tumor progression. (A) Spatial transcriptome mapping of fundus nodule biopsy sections collected from the same patient depicted in Figure 4 (A) at the same time. The diagnosis progressed from low-grade MALT to DLBCL in this subsequent biopsy. Left: sections from two different regions underwent 10μm-microfluidic device spatial barcoding. Right top: unsupervised clustering of Region 1 identified two clusters. Right bottom: unsupervised clustering of Region 2 revealed 10 transcriptomically distinct subpopulations. (B) Spatial characterization of representative cell types based on the expression of signature gene. Genes defining each module score are listed. (C) Spatial heterogeneities and interactions among tumor B cells. Left top: comparative analysis of chemokine gene expression between clusters 2 and 5. Left bottom: signaling pathways regulated by DEGs between cluster 2 vs. cluster 5. Right: spatial distribution of the Chemokine Score and RhoA Signaling Score. Genes defining each module score are listed. (D) Cellular-level spatial mapping unveils a distinct transcriptomic neighborhood. Left: comparative analysis of gastric mucus-secreting cell related gene expression between clusters 4, 7, and 8. Right top: enlarged transcriptomic neighborhood highlighted by white square in (A). Right bottom: tissue morphology of the corresponding area defined by H&E staining of an adjacent section. (E) Spatial analysis elucidates the molecular dynamics driving tumor progression. Left: schematic illustration showing comparative analysis. Right: signaling pathways regulated by DEGs between tumor B cells in DLBCL vs. MALT biopsy, revealing a significant upregulation of NF-κB signaling and its associated upstream and downstream pathways. (F) Expression comparison of key genes involved in the NF-κB signaling between DLBCL vs. MALT biopsy. (G) IHC staining for Ki67 on adjacent sections from the two biopsies. (H) Spatial expression mapping of genes encoding plasma cell kappa and lambda chains in the two biopsies. (I) ISH staining for kappa and lambda chain mRNA in the designated area in (H). (J) Distance distribution between macrophages and tumor B cells in the two biopsies. Significance level was calculated with two-tailed Mann-Whitney test, **** P < 0.0001. (K) Signaling pathways regulated by DEGs between macrophages in DLBCL vs. MALT biopsy, revealing a significant upregulation of macrophage alternative activation signaling and its associated pathways. (L) Ligand-receptor interactions between macrophage cluster 1 and tumor B cell clusters 2 and 5. The distinctive communication pattern of TGF-β (*TGFB1*) and the integrin family (*ITGB1*, *ITGB5*, and *ITGB8*) is indicated and spatially visualized. Edge thickness is proportional to correlation weights. In (C), (E) and (K), *z* score is computed and used to reflect the predicted activation level (*z*>0, activated; *z*<0, inhibited; *z*≥2 or *z*≤−2 can be considered significant).

Thoughtfully selected biopsies collected from different anatomical sites at the same timepoint enabled us to elucidate the spatial molecular dynamics propelling tumor progression. Comparative analysis of gene expression profiles in the tumor region, transitioning from low-grade MALT to high-grade DLBCL, revealed a significant upregulation of NF-κB signaling and its associated upstream and downstream pathways (Figure 7E). Furthermore, canonical tumor suppression pathways, including p53, p14/p19ARF, and PTEN, exhibited notable inhibition. We observed elevated expression of a range of key genes involved in the NF-κB signaling pathway, particularly *AKT1*, *IKBKB*, *PIK3C3*, and *PIK3R5* (Figure 7F). The considerable NF-κB activation, a hallmark of aggressive DLBCL^91^, could be associated with their significantly active c-Myc (MYC) signaling, spatially concentrated in the tumor B cell zone (Figure S7F). This was confirmed by pronounced c-Myc IHC staining presented in the DLBCL biopsy (Figure S7G). We also found a strong positive correlation between MYC signaling and miR-21 expression (Figure S7H), aligning with the documented role of c-Myc in activating miR-21 by directly binding to its promoter^92^. The coordinated upregulation of these signaling pathways, functional molecules, and their interactive networks may contribute to the high proliferative index in the DLBCL section, as indicated by its high Ki67 proliferative activity (∼70%) compared to the MALT section (<10%) (Figure 7G).

The detection of B-cell clonality has proven valuable for diagnosing B-cell lymphomas^93^. Abnormalities in light chain composition can lead to a significant elevation in one of the chains, resulting in an abnormal kappa:lambda ratio. In the MALT biopsy, a comparable expression level of the kappa chain constant domain gene *IGKC* and lambda chain genes, including *IGLC1*, *IGLC2*, and *IGLC3*, was observed and confirmed by ISH stain for kappa and lambda mRNA (Figure 7H). The finding of polytypic light chains in this biopsy is likely due to the indolent stage, where no clone dominates yet and relatively less plasmacytic differentiation. However, in the progressed DLBCL biopsy, the plasma cells were mostly kappa type, represented by a dominant expression of *IGKC*, which could be associated with greater differentiation status. Consistently, we observed a significant upregulation of pathways linking inflammation to cancer and involved in tumor cell survival and malignant progression, alongside a downregulation of principal tumor suppression pathways in the DLBCL plasma cells compared to those in the MALT counterpart (Figures S7I and S7J). Notably, this patient had been followed for lymphadenopathy until the identification of enlarged lymph nodes 3 years later. This led to biopsy of a 6 cm neck lymph node that revealed a recurrence of DLBCL with kappa-restricted monoclonal tumor B cells, providing robust evidence for the detection accuracy and potential diagnostic value of Patho-DBiT.

Finally, we investigated the functional profile of macrophages and their spatial interaction with tumor cells in the two biopsies. In the MALT lymphoma section, macrophages in cluster E4 remained distantly positioned from the tumor zone in cluster E1 (Figure 4A). In contrast, a profound infiltration of macrophages (cluster 1) into the tumor region (clusters 2/5) was observed in the DLBCL section (Figure 7A), resulting in a significantly reduced macrophage-tumor distance in the latter (Figure 7J). The DEGs between macrophages in DLBCL and MALT lymphoma regulated a significant activation of the macrophage alternative activation signaling pathway (Figures 7K and S7K). Additionally, associated pathways involved in intercellular communications, metabolic modulations, and intracellular responses were affected, potentially contributing to the roles of tumor-associated macrophages in promoting tumor growth and invasion^94^. The deeper exploration of ligand-receptor interactions shed light on the communication network between macrophages and B cells. This analysis unveiled a plethora of crucial pairs activated within the DLBCL microenvironment (Figure 7L), highlighting the potential role of *APOE*-*LRP1* pair in metabolic shaping of the conversion from classic (M1) to alternative (M2) macrophage^95^, CXCL9-CXCR3 axis in orchestrating the recruitment of effector T cells^96^, and VEGFC-IL6ST pair in fostering tumor lymphangiogenesis^97^. A noteworthy interaction was the crosstalk between TGF-β (*TGFB1*) and the integrin family (*ITGB1*, *ITGB5*, and *ITGB8*), potentially instigating the formation of pro-tumorigenic M2-macrophages and facilitating tumor immune escape^98^. The spatial expression pattern of this interaction can be distinctly visualized. Collectively, our high-sensitivity Patho-DBiT empowers us to spatially map the molecular evolution from low-grade to high-grade tumor at cellular level resolution, deepening our understanding of the complex interplay shaping the tumor microenvironment in transformed large B-cell lymphomas.

## Discussion

Despite recent breakthroughs in spatial biology, either imaging-based or sequencing-based spatial transcriptomics is largely equivalent to spatial gene expression quantification. However, an RNA molecule experiences a complex life cycle involving transcription, splicing, maturation, translation, and degradation^99,100^. It is highly desirable to profile all these RNA species spanning their life cycle to explore the biology of RNA at genome scale and cellular level. Furthermore, FFPE is the most widely used method for banking clinical tissue biospecimens. The ability to explore such rich information in archival FFPE tissue specimens is poised to transform the study of human disease biology and clinical histopathology. Herein, we developed Patho-DBiT, a first-of-its-kind technology enabling the spatial exploration of rich RNA biology in FFPE tissues. Counter-intuitively, a variety of RNA species including small RNA molecules, although highly fragmented, are well preserved in the FFPE blocks. Combining polyadenylation and spatial deterministic barcoding enabled high-yield detection of these molecules with genome-wide coverage and excellent depth. Thus, this unique property has led to enhanced coverage of mature mRNA transcripts and facilitates the capture of small noncoding RNAs lacking poly(A) tails. We demonstrated this ability by spatially profiling the expression of microRNAs. Patho-DBiT not only aligns reads to pre-microRNA regions but also accurately demarcate the boundaries of known dominant mature microRNA strands produced from the pre-microRNA hairpins. These examples include hsa-miR-143-3p, hsa-miR-145-5p (Figure 5C), hsa-miR-155-5p (Figure 5G), hsa-miR-30b-5p, hsa-miR-30d-5p, hsa-miR-30e-5p (Figure S5A), hsa-miR-146a-5p, hsa-miR-150-5p (Figure S5B), and mmu-miR-122-5p (Figure S1H). Remarkably, unlikely most other microRNA genes, hsa-miR-142 precursor yields both 5p and 3p mature microRNAs at high levels, and this feature was faithfully captured by Patho-DBiT (Figure S5B). The reason why small RNAs such as microRNAs are well preserved to allow the detection by Patho-DBiT, even in aged FFPE blocks, is not fully clear, but may be due to the protection by RNA-binding proteins such as the Argonaute family that bind to mature microRNAs. The ability of the Patho-DBiT data to retrieve known regulatory relationships between microRNAs and cellular pathways raises the possibility to use this technology in the future to discover important gene regulation mechanisms in disease settings.

Beyond profiling gene expression, Patho-DBiT enables spatial profiling of post-transcriptional RNA processing, including alternative splicing and A-to-I RNA editing. Compared to other spatial transcriptomics methods, Patho-DBiT achieved substantially broader coverage across the gene body (Figure S2C), allowing for transcript isoform analysis with heightened sensitivity (Figure S2D When applied to an FFPE mouse coronal brain section, Patho-DBiT uncovers hundreds of differential alternative splicing events between individual pairs of brain regions (Figure 2E), particularly in genes with vital brain functions (Figures 2F, 2G, and 2H). Furthermore, Patho-DBiT identifies regionally variable RNA editing events (Figure 2I) that correlate with the regional variations in the gene expression level of *Adarb1*, responsible for encoding the brain-expressed RNA editing enzyme (Figure 2J). Collectively, Patho-DBiT opens new avenues for exploring the spatial patterns and regulation of RNA processing within FFPE tissues. As illustrated by the example of *Adarb1*, correlating the spatial expression profiles of RNA binding proteins with the spatial distributions of transcript isoforms may unveil novel regulators of spatially resolved RNA processing events.

Pathology is a cornerstone in cancer diagnosis and treatment. It has evolved significantly with technological advances, moving from microscopic observation of tissues to sophisticated genomic analysis. By strategically appending poly(A) tails to RNAs, fragmented during the embedding process and further degraded due to long-term storage at room temperature, Patho-DBiT enabled high-sensitivity spatial transcriptomics profiling of a 5-year archival AITL sample (Figure 3). Boasting a remarkable detection of 5,364 genes and 11,989 UMIs per 50μm pixel, our innovative platform even outshines the current gene capture efficiency in fresh frozen tissues, facilitating a faithful reconstruction of spatial cellular compositions and empowering the identification of AITL-specific molecular pathways that drive lymphomagenesis. In the characterization of a 3-year archival MALT tissue, Patho-DBiT showcased its prowess in extracting a wealth of information from a single run, ranging from exonic mRNA expression, genome-scale single nucleotide variants, microRNA-mRNA regulatory networks, and splicing dynamics linked to developmental trajectories, with rigorous validation of accuracies using traditional staining methods including H&E, IHC, IF, and ISH (Figures 4-6). The result is a finely crafted spatial molecular landscape that serves as a powerful resource for pathologists to disentangle the intricate biological process underlying tumorigenesis.

Spatial structure of tumor profoundly influences disease development and response to treatment. To propel personalized medicine to the next level, there is a critical imperative to spatially evaluate tumor microenvironment at cellular level and identify target candidates within the dynamically evolving and heterogeneous neoplastic population. As demonstrated in this study of indolent MALT and DLBCL cases, the same patient can harbor similar B-cell tumor cells that take advantage of different microenvironments to survive and proliferate (Figure 7). Patho-DBiT holds the potential to revolutionize the analysis of human pathology by revealing vulnerabilities in the structural supports of tumors and providing real-time insights into molecular kinetics. Furthermore, as tumors persistently develop resistance to targeted therapies, including immunotherapies^101^, we envision that this technology can assist clinicians in identifying future trajectories of the tumor and responding promptly with synergistic compounds. Nevertheless, in order to harness emerging genomic knowledge and bring it to bear on current pathology diagnostics, newly developed tools like Patho-DBiT must undergo thorough testing and validation on patient samples, keeping in mind that the traditional skills of clinical observation and recognition of spatial patterns may still offer valuable lessons in this context.

## Acknowledgement

We extend our appreciation to Dr. Lei Wang from Yale West Campus cleanroom for assistance in fabricating the microfluidic master wafers, the pathology team for their efforts in collecting patient specimens and embedding tissues in FFPE blocks, and the YPTS team for their contributions to FFPE tissue sectioning and staining. Computational data analysis was conducted with the Yale High Performance Computing clusters (HPC). We acknowledge the support received from the U.S. National Institutes of Health including grants RF1MH128876, U54AG079759, UH3CA257393, U54AG076043, U54CA274509, R01CA245313, RM1MH132648 (all to R.F.), R33CA246711 (to R.F. and J.L.), the support from the Packard Fellowship for Science and Engineering 2012-38215 (to R.F.), and additional NIH grants: R01GM138856 (to J.L.), U54DK106857 (to J.L.), R56HG012310 (to Y.X.), U54CA268083 (subaward to R.F.), UM1MH130991 (subaward to R.F.).

## Author Contributions

Conceptualization, R.F., Z.B., M.L.X., J.L., and Y.X.; data curation, Z.B., D.Z., and Y.G.; formal analysis, Z.B., D.Z., Y.G., B.T., Daiwei.Z., G.S., M.Y., X.T., M.G., and M.L.; funding acquisition, R.F., M.L.X., J.L., and Y.X.; investigation, Z.B., S.B., A.E., B.T., and N.Z.; methodology, Z.B., R.F., J.L., and B.T.; project administration, Z.B. and R.F.; resources, M.L.X., Z.B., A.E., and S.B.; software, D.Z. and Y.G.; scientific discussion, Yang.X. and Y.L.; writing–original draft, Z.B. and B.T. with inputs from D.Z., Y.G., S.B., and A.E.; writing–review and editing, R.F., M.L.X, J.L., and Y.X.

## Methods

### Patient specimens

De-identified archival formalin-fixed paraffin-embedded (FFPE) human lymphoma tissue blocks, originally collected by physicians for diagnostic purposes, were sourced from the Yale Pathology Tissue Services (YPTS), a Pathology-based Central Tissue Resource Lab that provides comprehensive tissue-related services and materials in de-identified format for investigators at Yale University. The tissue collection was conducted with Yale University Institutional Review Board approval with oversight by Tissue Resource Oversight Committee. Written informed consent for participation, including cases where identification was collected alongside the specimen, was obtained from patients or their guardians, adhering to the principles of the Declaration of Helsinki. Each sample was handled in strict compliance with HIPAA regulations, University Research Policies, Pathology Department diagnostic requirements, and Hospital by-laws. The excisional biopsy from the left upper arm subcutaneous nodule was collected and embedded in 2018 from a patient presenting with angioimmunoblastic T-cell lymphoma (AITL) in multiple lymph nodes and subcutaneous sites. Biopsies from the gastric antrum revealing marginal zone lymphoma of mucosa-associated lymphoid tissue (MALT) and the fundus nodule indicating diffuse large B-cell lymphoma (DLBCL) were collected and embedded in 2020. These biopsies were obtained from a patient who incidentally presented with retroperitoneal lymphadenopathy during imaging originally performed for an orthopedic visit. Upper endoscopy revealed multiple areas of erosion in the stomach, and a breath test for H. pylori was positive.

### Surgical pathology report of the lymphoma biopsies

The AITL sections show a sheet of lymphocytes, some with atypical morphology. There are thick and thin bands of fibrosis and interspersed blood vessels. The atypical cells have irregular to round nuclei, speckled chromatin, variable small nucleoli, and a small amount of cytoplasm. There is infiltration into the adjacent fat. Significant mitotic figures, apoptotic figures, or necrosis is not identified. The atypical cells are CD3-positive T cells that are positive for CD4, CD2, CD5, CD10, CXCL13, and PD-1. They are negative for CD25 and CD8 with partial loss of CD7. There are abundant background CD20-postive B cells. The Ki-67 proliferation index is overall approximately 20-30%. T cell gene rearrangement was positive, showing same peaks as other sites of involvement. Flow cytometric analysis reveals CD4+ T cells are increased in the specimen, representing about 36% of total lymphocytes with few CD8+ elements detected. In addition, CD4+ T cells possess an abnormal immunophenotype.

The MALT sections reveal gastric antral mucosa with numerous lymphoid follicles showing monotonous small lymphocytes that demonstrate ovoid nuclei, condensed chromatin, and indistinct nucleoli. No large cell component is seen in this part. The tumor cells are CD20 positive B cells that co-express BCL-2 and CD43, are negative for CD5, CD10, BCL-6, CD23, LEF1, and cyclinD1. Ki-67 is low at <10%. CD3 highlights scattered small T cells. H.pylori immunostaining is negative.

The DLBCL sections reveal sheets of large pleomorphic lymphocytes, some with horseshoe shaped nuclei, dispersed chromatin, prominent nucleoli, and moderate amount of eosinophilic cytoplasm. There are numerous eosinophils in the background and no substantial small cell lymphoma. The tumor cells are positive for CD20, CD43, and MUM1 and negative for CD10, cyclinD1, and CD30. BCL-6 is faintly expressed in <20% of cells. C-myc is expressed in >80% of tumor cells and BCL-2 is expressed in >70% of cells. Ki-67 proliferation index is approximately 70%. CD3 positive small T cells are scattered. Para-aortic lymph node biopsy performed simultaneously shows involvement by metastatic DLBCL.

### Mouse paraffin tissues

The mouse E13 embryo, caudal hippocampus coronal brain/Region.9, and lymph node sections were purchased from Zyagen (San Diego, CA). Tissues were freshly harvested from C57BL/6 mice fixed in 10% Neutral Buffered formalin and processed for embedding in low temperature melting paraffin. All tissue preparation steps from harvesting to embedding in paraffin were done in RNase-, DNase-, and protease-free conditions. Tissue sections were hematoxylin and eosin (H&E) stained and examined by histologists with extensive experience to be sure of excellent morphology and high quality.

### Sample handling and section preparation

For both human and mouse samples, paraffin blocks were sectioned at a thickness of 7-10 µm and mounted on the center of Poly-L-Lysine coated 1 x 3” glass slides. Serial tissue sections were collected simultaneously for Patho-DBiT and other staining. The sectioning of lymphoma patient samples was carried out at YPTS, while mouse sectioning was performed by Zyagen technicians. Paraffin sections were shipped in tightly closed slide boxes or slide mailers at room temperature and stored at −80°C upon receipt until use.

### Fabrication of microfluidic device

The comprehensive fabrication process, employing standard soft lithography, has been detailed in our previous publication^13^. Briefly, high-resolution chrome photomasks with a customized pattern were printed and ordered from Front Range Photomasks (Lake Havasu City, AZ). Upon receipt, the masks underwent cleaning with acetone to remove any dirt or dust. Master wafers were then produced using SU-8 negative photoresist (SU-2010 or SU-2025) on silicon wafers following the manufacturer’s guidelines, with feature width of 50 µm, 20 µm, or 10 µm. The newly fabricated wafers were treated with chlorotrimethylsilane for 20 minutes to develop high-fidelity hydrophobic surfaces. Subsequently, polydimethylsiloxane (PDMS) microfluidic chips were fabricated through a replication molding process. The base and curing agents were mixed thoroughly with a 10:1 ratio following the manufacturer’s guidelines and poured over the master wafers. After degassing in the vacuum for 30 min, the PDMS was cured at 70°C for at least 2 hours. The solidified PDMS slab was cut out, and the inlets and outlets were punched for further use.

### DNA barcodes annealing

DNA oligos used in this study were procured from Integrated DNA Technologies (IDT, Coralville, IA) and the sequences were listed in Table S2. Barcode (100 µM) and ligation linker (100 µM) were annealed at a 1:1 ratio in 2X annealing buffer (20 mM Tris-HCl pH 8.0, 100 mM NaCl, 2 mM EDTA) with the following PCR program: 95°C for 5 min, slow cooling to 20°C at a rate of −0.1°C/s, followed by 12°C for 3 min. The annealed barcodes can be stored at −20°C until use.

### Tissue deparaffinization and decrosslinking

Tissue section was retrieved from the −80°C freezer and equilibrated to room temperature for 10 minutes until all moisture dissipated. Following this, the tissue slide underwent a 1-hour baking process at 60°C to facilitate softening and melting of the paraffin. Removal of paraffin was achieved by immersing slides in Xylene for two changes, followed by rehydration in a series of ethanol dilutions, including two rounds of 100% ethanol and once each of 90%, 70%, and 50% ethanol, culminating in a final wash with distilled water. Each step was performed for a duration of 5 min. Subsequently, the tissue slide was submerged in 1X antigen retrieval buffer and subjected to steaming using boiling water for 30 minutes, followed by a 30-minute cooldown to room temperature. After a brief dip in distilled water, intact tissue scan was captured using a 10X objective on the EVOS M7000 Imaging System.

### Permeabilization, *in situ* polyadenylation, and reverse transcription

The tissue was permeabilized for 20 min at room temperature with 1% Triton X-100 in DPBS, followed by 0.5X DPBS-RI (1X DPBS diluted with nuclease-free water, 0.05 U/µL RNase Inhibitor) wash to halt permeabilization. The tissue slide was then air-dried and equipped with a PDMS reservoir covering the region of interest (ROI). In situ polyadenylation was performed using *E. coli* Poly(A) Polymerase. Initially, samples were equilibrated by adding 100 µL wash buffer (88 µL nuclease-free water, 10 µL 10X Poly(A) Reaction Buffer, 2 µL 40 U/µL RNase Inhibitor) and incubating at room temperature for 5 min. Following wash buffer removal, 60 µL of the Poly(A) enzymatic mix (38.4 µL nuclease-free water, 6 µL 10X Poly(A) Reaction Buffer, 6 µL 5U/µL Poly(A) Polymerase, 6 µL 10mM ATP, 2.4 µL 20 U/µL SUPERase•In RNase Inhibitor, 1.2 µL 40 U/µL RNase Inhibitor) was added to the reaction chamber and incubated in a humidified box at 37°C for 30 min. To remove excessive reagents, the slide was dipped in 50 mL DPBS and shake-washed for 5 min after the reaction. Subsequently, 60 µL of the reverse transcription mix (20 µL 25 µM RT Primer, 16.3 µL 0.5X DPBS-RI, 12 µL 5X RT Buffer, 6 µL 200U/µL Maxima H Minus Reverse Transcriptase, 4.5 µL 10mM dNTPs, 0.8 µL 20 U/µL SUPERase•In RNase Inhibitor, 0.4 µL 40 U/µL RNase Inhibitor) was loaded into the PDMS reservoir and sealed with parafilm. The sample was incubated at room temperature for 30 min and then at 42°C for 90 min, followed by a 50 mL DPBS wash as described before.

### Spatial barcoding with microfluidic devices

To ligate barcode A in situ, the first PDMS device was meticulously positioned atop the tissue slide, aligning the 50 center channels over the ROI. The chip was imaged to record the positions for downstream alignment and analysis. Afterwards, an acrylic clamp was applied to firmly secure the PDMS to the slide, preventing any inter-channel leakage. The ligation mix, comprising 100 µL 1X NEBuffer 3.1, 61.3 µL nuclease-free water, 26 µL 10X T4 ligase buffer, 15 µL T4 DNA ligase, 5 µL 5% Triton X-100, 2 µL 40 U/µL RNase Inhibitor, and 0.7 µL 20 U/µL SUPERase•In RNase Inhibitor, was then prepared. For the barcoding reaction, 5 µL of the ligation solution, containing 4 µL ligation mix and 1 µL 25 µM DNA barcode A (A1-A50), was introduced into each of the 50 inlets. The solution was withdrawn to flow through the entire channel using a delicately adjusted vacuum. After a 30-minute incubation at 37°C, the PDMS chip was removed, and the slide was washed with 50 mL DPBS. Subsequently, the second PDMS device, featuring 50 channels perpendicular to the first PDMS, was attached to the ROI on the air-dried slide. A bright-field image was captured, and the ligation of barcode B set was performed similarly. Finally, after five flow-washes with 1 mL nuclease-free water to remove residual salt, the final scan was conducted to record the microchannel marks imprinted onto the tissue ROI.

### Tissue lysis and cDNA extraction

The barcoded tissue ROI was enclosed with a clean PDMS reservoir and securely clamped using acrylic chips. A 2X lysis buffer was prepared in advance, consisting of 20 mM Tris-HCl pH 8.0, 400 mM NaCl, 100 mM EDTA, and 4.4% SDS. For tissue digestion, 70 µL of the lysis mix (30 µL 1X DPBS, 30 µL 2X lysis buffer, 10 µL 20 µg/µL Proteinase K solution) was loaded into the PDMS reservoir, sealed with parafilm, and incubated in a humidified box at 55°C for 2 hours. After the reaction, the parafilm was removed, and all the liquid containing cDNA was collected into a 1.5mL DNA low-bind tube. Additionally, 40 µL of fresh lysis mix was loaded into the reservoir to collect any remaining cDNA material. The tissue lysate was incubated overnight at 55°C to completely reverse crosslinks, after which it could be stored at −80°C until the subsequent steps.

### cDNA purification, template switch, and PCR amplification

To inhibit Proteinase K activity, 5 µL of 100 µM phenylmethylsulfonyl fluoride (PMSF) in ethanol was introduced into the lysate and incubated at room temperature for 10 min with rotation. Following this, ∼35 µL of nuclease-free water was added to adjust the total volume to 150 µL. The cDNA was purified using 40 µL of Dynabeads MyOne Streptavidin C1 beads resuspended in 150 µL of 2X B&W buffer (10 mM Tris-HCl pH 7.5, 1 mM EDTA, 2 M NaCl). The mixture was incubated at room temperature for 60 min with rotation to ensure sufficient binding, followed by magnetic separation and two washes with 1X B&W buffer with 0.05% Tween-20, and an additional two washes with 10 mM Tris-HCl pH 7.5 containing 0.1% Tween-20. Streptavidin beads bound with cDNA molecules were then resuspended in 200 µL of TSO Mix (75 µL nuclease-free water, 40 µL 5X RT buffer, 40 µL 20% Ficoll PM-400, 20 µL 10mM dNTPs, 10 µL 200U/µL Maxima H Minus Reverse Transcriptase, 5 µL 40 U/µL RNase Inhibitor, 10 µL 100 µM TSO Primer). The template switch reaction was conducted at room temperature for 30 min and then at 42°C for 90 min with gentle rotation. After a single wash with 10 mM Tris-HCl pH 7.5 containing 0.1% Tween-20 and another wash with nuclease-free water, the beads were resuspended in 200 µL of PCR Mix (100 µL 2X KAPA HiFi HotStart ReadyMix, 84 µL nuclease-free water, 8 µL 10 µM PCR Primer 1, 8 µL 10 µM PCR Primer 2). This suspension was then distributed into PCR stripe tubes. An initial amplification was conducted with the following PCR program: 95°C for 3 min, cycling five times at 98°C for 20 s, 63°C for 45 s, 72°C for 3 min, followed by an extension at 72°C for 3 min and 4°C hold. Following magnetic removal of the beads, 19 µL of the PCR solution was combined with 1 µL 20X EvaGreen for quantitative real-time PCR (qPCR) analysis using the same program. The remaining samples underwent further amplification, with the cycle numbers determined by 1/2 of the saturated signal observed in qPCR results. The PCR product was then purified using SPRIselect beads at a 0.8X ratio, adhering to the standard manufacturer’s instructions. The resulting cDNA amplicon underwent analysis using a TapeStation system with D5000 DNA ScreenTape and reagents. This stage provides a secure stopping point, allowing the sample to be stored at −20°C until the next steps.

### rRNA removal, library preparation, and sequencing

The SEQuoia RiboDepletion Kit was employed to eliminate fragments derived from rRNA and mitochondrial rRNA from the amplified cDNA product, following the manufacturer’s guidelines. Based on the TapeStation readout profile, 20 ng of cDNA was used as the input amount, and three rounds of depletion were performed. Subsequently, 7 cycles of the aforementioned PCR program were executed to directly ligate sequencing primers, using a 100 µL system consisting of 50 µL 2X KAPA HiFi HotStart ReadyMix, ∼42 µL solution from the rRNA removal step, 4 µL 10 µM P5 Primer, and 4 µL 10 µM P7 Primer. The resulting library underwent purification using SPRIselect beads at a 0.8X ratio, quality control checked using TapeStation, and was then sequenced on an Illumina NovaSeq 6000 Sequencing System with a paired-end 150bp read length.

### CODEX spatial phenotyping using PhenoCycler-Fusion

Spatial high-plex phenotyping of the adjacent FFPE section was performed following the CODEX PhenoCycler-Fusion user guide (https://www.akoyabio.com/wp-content/uploads/2021/01/CODEX-User-Manual.pdf). Briefly, the tissue section underwent deparaffinization, hydration, antigen retrieval and equilibration in staining buffer, followed by antibody cocktail staining incubated at room temperature for 3 hours in a humidity chamber. After the completion of the incubation, a series of sequential steps, including post-fixation, ice-cold methanol incubation, and a final fixative step, were performed. The tissue section, attached to the flow cell, was then incubated in 1X PhenoCycler buffer with additive for a minimum of 10 minutes to enhance adhesion. Afterwards, the CODEX cycles were configured, the reporter plate was prepared and loaded, and the imaging process commenced. Upon completion of the imaging cycles, a final QPTIFF file was generated, which could be visualized using QuPath V0.5.0^102^. Information about PhenoCycler antibody panels, experimental cycle design, and reporter plate volumes can be found in Table S3.

### H&E, immunohistochemistry (IHC) and in situ hybridization (ISH)

Histological H&E staining and clinical-level IHC and ISH on adjacent FFPE sections were conducted at Yale University School of Medicine, Department of Pathology and at YPTS. These procedures adhered to Clinical Laboratory Improvement Amendments (CLIA)-certified laboratory protocols as well as YPTS’s rigorous standard protocols, ensuring precision and accuracy in the analysis of tissue samples.

### Immunofluorescence staining (IF)

The adjacent FFPE sections underwent a standard IF procedure. After deparaffinization and antigen retrieval, the tissue sections were fixed in 4% formaldehyde for 10 minutes and subsequently blocked with DPBS containing 5% bovine serum albumin for 1 hour at room temperature. CD68 antibodies, diluted at 1:100 in the blocking buffer, were applied and left to incubate overnight at 4°C. Secondary antibodies for CD68, Alexa-594 labeled CD138, and Alexa-647 labeled CD20 were then introduced following a standard IF protocol, with a 30-minute incubation at room temperature. The nuclei were counterstained with DAPI at a 1:4000 dilution. Imaging was conducted using a Leica TCS SP5 Confocal microscope.

### Sequence alignment and generation of gene expression matrix

To decode sequencing data, the FASTQ file Read 2 underwent processing, involving the extraction of unique molecular identifiers (UMIs) and spatial Barcode A and Barcode B. The Read 1 containing cDNA sequences was trimmed using Cutadapt V3.4^103^ and then aligned to either the mouse GRCm38-mm10 or human GRCh38 reference genome using STAR V2.7.7a^104^. Utilizing ST_Pipeline V1.7.6^105^, spatial barcode sequences were demultiplexed based on the predefined coordinates of the microfluidic channels and ENSEMBL IDs were converted to gene names, generating the gene-by-pixel expression matrix for downstream analysis. Matrix entries corresponding to pixel positions devoid of tissues were excluded.

### Gene data normalization and unsupervised clustering analysis

Spatial gene expression analysis was conducted through the Seurat V4 pipeline^106^. First, SCTransform, designed for normalization and variance stabilization in single-cell RNA sequencing (scRNA-seq) datasets^107^, was employed to normalize gene expression within each pixel. Linear dimensional reduction was performed using the “RunPCA” function, and the optimal number of principal components for subsequent analysis was determined through a heuristic method, generating an ‘Elbow plot’ that ranks PCA components based on their percentage of variances. Second, “FindNeighbors” function was utilized to embed pixels in a K-nearest neighbor graph structure based on the Euclidean distance in PCA space, and “FindClusters” was implemented using a modularity optimization technique to cluster the pixels. Finally, the non-linear dimensional reduction function “RunUMAP” was applied to visually explore spatial heterogeneities using the Uniform Manifold Approximation and Projection (UMAP) algorithm, and the identification of differentially expressed genes (DEGs) defining each cluster was accomplished through the “FindMarkers” function for pairwise comparison between groups of pixels.

### Integration with scRNA-seq datasets

At a resolution of 50 µm, Patho-DBiT assay pixels capture the expression profiles of multiple cells. We employed the ‘anchor’-based integration workflow included into Seurat V4 to deconvolute each spatial voxel, predicting the underlying composition of cell types^108^. This facilitated the probabilistic transfer of annotations from a reference to a query set. After standard “SCTransform” normalization of both Patho-DBiT and reference scRNA-seq data, the “FindTransferAnchors” function identified anchors between the reference scRNA-seq and our query Patho-DBiT object. Subsequently, the “TransferData” function was applied for label transfer, providing a probabilistic classification for each spatial pixel based on well-annotated scRNA-seq identities. These predictions were added as a new assay to the Patho-DBiT object. Unsupervised clustering was then performed on the combined Patho-DBiT and reference dataset, resulting in an integrated UMAP where Patho-DBiT pixels were projected onto the scRNA-seq cluster landscape. The mouse organogenesis reference dataset was obtained from GSE119945^14^, and the mouse brain cortex and hippocampus reference dataset was downloaded from the Allen Mouse Brain Atlas (https://portal.brain-map.org/atlases-and-data/rnaseq)^26^.

### qPCR analysis of rRNA removal efficiency

To assess the rRNA removal efficiency of Patho-DBiT, qPCR analysis was performed on cDNA amplicons obtained from three independent FFPE mouse E13 embryos before and after rRNA removal. Each sample, with an input amount of 2.5 ng cDNA, underwent a total volume of 25 µL in the KAPA HiFi HotStart ReadyMix reaction system. Forward and reverse primers targeting cytoplasmic (5S, 5.8S, 18S, and 28S) and mitochondrial (12S and 16S) rRNA were custom-designed and ordered from IDT (sequences provided in Table S4). QuantiTect Primer Assays for mouse GAPDH and β-actin genes served as internal controls. The qPCR reactions were conducted on a CFX Connect Real-Time System, and fold changes were determined using the comparative CT method^109^.

### Gene body coverage calculation

For each sample, we computed the percentile coverage along the gene body from 5’ to 3’ using the “geneBody_coverage.py” module from the RSeQC package V5.0.1 with default settings^110^.

### Spatial alternative splicing analysis

For both our Patho-DBiT FFPE mouse brain and 10x Genomics Visium FFPE or fresh-frozen mouse brain samples, we evaluated five types of alternative splicing events (SE, RI, A3SS, A5SS, MXE) and their respective splice-junction-spanning read counts from the Binary Alignment Map (BAM) file of each sample. The rMATS-turbo pipeline V4.1.2^28^ with parameters “-t single --allow-clipping --variable-read-length” and the GRCm38-mm10 mouse gene annotation were employed for this analysis. Candidate events were considered for further analysis if their inclusion and skipping isoform read counts were both ≥ 2 when aggregated from all pixels within the sample. Within each spatial pixel, a gene was deemed to have alternative splicing information if at least one splice-junction-spanning read of either inclusion or skipping isoform was detected. To identify alternative splicing events showing regional differences in our Patho-DBiT data, pseudo-bulk BAM files of each brain region were generated by merging reads from all the pixels within the same region. Pairwise regional differential alternative splicing analysis was performed by running rMATS-turbo on the generated pseudo-bulk BAM files for each pair of two regions. An alternative splicing event was considered significant if it exhibited an exon inclusion level difference of > 0.05 between two regions, with a false discovery rate (FDR) of ≤ 0.05. Exon inclusion levels and FDRs were obtained from rMATS-turbo’s splice-junction-read-based outputs (*.MATS.JC.txt). The spatial locations of reads corresponding to alternative splicing events were deciphered using their barcode sequences, resulting in distinct inclusion and skipping isoform expression matrices for each event type. Seurat V4’s “NormalizeData” with a “LogNormalize”-based global-scaling normalization was applied, and the “SpatialFeaturePlot” was employed to visualize the spatial distribution of selected isoforms. The download links for the 10x Genomics datasets can be found in Table S5.

### Spatial adenosine-to-inosine (A-to-I) RNA editing analysis

A total of 107,095 reference mouse A-to-I RNA editing sites were retrieved from the REDIportal database (http://srv00.recas.ba.infn.it/atlas/search_mm.html)^111^ as of the download date on 9-20-2023. The counts of edited and unedited reads for each editing site were calculated from the BAM file containing all spatial pixels using the “mpileup” subcommand of samtools V1.16.1^112^, with parameters “--no-output-ins --no-output-ins --no-output-del --no-output-del --no-output-ends -B -d 0 -Q 25 -q 25” along with the reference editing site list and the GRCm38-mm10 mouse reference genome. Reads with bases “A” and “G” at editing sites were classified as unedited and edited, respectively. Candidate A-to-I RNA editing sites for further analysis were defined as those with a total coverage of ≥ 10 and an edited read count of ≥ 1 when aggregated from all pixels within the sample. The overall editing ratio for each editing site was computed by dividing the total number of edited reads across all pixels by the total coverage of that site. Similarly, the average editing ratio for each pixel or brain region was determined by dividing the total edited reads by the total coverage of all editing sites within that specific area. The reference spatial dataset, containing editing sites, editing ratios, and total read counts from long-read Nanopore sequencing of fresh frozen mouse brain sections, was obtained from the literature^42^. For comparison with our Patho-DBiT dataset, only sites with ≥10 long reads were included.

### Spatial microRNA alignment and analysis

The transcriptome output function of STAR was used to generate the microRNA transcriptome BAM file using annotations obtained from miRBase^113^. Only primary alignment of each read mapped to microRNA was preserved, and microRNAs with detected UMI count ≥1 were included in the downstream analysis. The nucleotide length of each mapped microRNA read was calculated and the count distribution across all identified microRNAs was generated. To visualize read coverage across the reference genomic region, the BAM file of specific microRNAs was directly imported into the Integrative Genomics Viewer (IGV)^114^, focusing on the precursor microRNA region, including the mature 5p-strand and 3p-strand, for detailed visualization. The spatial microRNA-by-pixel expression matrix was generated by decoding barcode sequences, and standard functions integrated into Seurat V4 were utilized for normalization and spatial visualization.

### Spatial single nucleotide variant (SNV) analysis

The germline variant calling pipeline, Strelka V2.9.10^115^, was utilized to identify potential SNVs from the mapped BAM file. Only high-confidence variant loci marked as “PASS” in Strelka, along with SNV sites having sequencing counts ≥60, were retained for further analysis. Each pixel and SNV site were assigned values: 0 for wild type, 1 for heterozygous mutation, or 2 for homozygous mutation. Positions with no detected mutated nucleotides were labeled as wild type, those with both mutated and wild-type nucleotides were classified as heterozygous mutation, and sites with only mutated nucleotides were categorized as homozygous mutation. For each pixel, only SNV sites identified by Strelka were incorporated into the profile, considering that RNA-seq data may not cover the entire genome^116^. By combining spatial coordinates defined by barcode sequences, a mutation-by-pixel matrix was generated, and the cumulative number of SNVs within each pixel was calculated to delineate spatial mutational burden. Subsequently, this mutational matrix was input into the Seurat V4 pipeline to perform unsupervised clustering analysis using standard normalization, dimensional reduction, and spatial visualization methods.

### Coverage comparison with 10x Genomics datasets

To assess and compare the genomic location coverage bandwidth between Patho-DBiT and 10x Genomics 3’ scRNA-seq or Visium spatial datasets, we obtained the aligned BAM files from the respective website. The sequencing depth was normalized by randomly selecting an equivalent number of reads in each 10x Genomics file and our Patho-DBiT data. Genomic regions with at least one detected read were considered covered. The download links for the 10x Genomics datasets can be found in Table S5.

### Spatial RNA splicing dynamics

The analysis involved extracting counts of spliced and unspliced reads independently from the aligned BAM file. Genomic regions corresponding to exons and introns were obtained from the GENCODE annotation^51^. Utilizing the “intersect” tool within bedtools V2.31.0^117^, reads overlapping with intronic regions were identified, and the associations between each read and its corresponding gene were documented. The remaining reads that overlapped with exonic regions were selected, and their connections to the overlapped genes were documented as well. After demultiplexing their spatial coordinates, reads containing region records were processed to generate spliced and unspliced count matrices, respectively. Following this, the two matrices were imported into the scVelo pipeline^71^, where RNA velocity, pseudotime analysis, and visualization were implemented using default settings. Pixel annotations, featuring assigned cluster identities, were transferred from the Seurat clustering analysis conducted on the combined exonic and intronic expression matrices.

### Ligand-receptor interaction analysis

The R toolkit Connectome V1.0.0^118^ was employed to investigate cell-cell connectivity patterns using ligand and receptor expressions from our Patho-DBiT datasets. The normalized Seurat object served as input, and cluster identities were utilized to define nodes in the interaction networks, resulting in an edgelist connecting pairs of nodes through specific ligand-receptor mechanisms. We selected top-ranked interaction pairs, prioritizing those more likely to be biologically and statistically significant based on the scaled weights of each pair. The “sources.include” and “targets.include” parameters were applied to specify the source cluster emitting ligand signals and the target cluster expressing receptor genes that sense the ligands.

### Ingenuity Pathway Analysis

Ingenuity Pathway Analysis (IPA, QIAGEN)^67^ was employed to uncover the underlying signaling pathways regulated by the DEGs characterizing each identified cluster or two groups. The DEG list, along with the corresponding fold change value, p-value, and adjusted p-value of each gene, was imported into the software. The Ingenuity Knowledge Base (genes only) served as the reference set for performing Core Expression Analysis. The z-score was utilized to assess the activation or inhibition level of specific pathways. Conceptually, the z-score is a statistical measure gauging how closely the actual expression pattern of molecules in our DEG dataset aligns with the expected pattern based on the literature for a particular annotation. A z-score >0 signifies activation or upregulation, while a z-score <0 indicates inhibition or downregulation. A z-score ≥2 or ≤−2 is considered significant. The p-value for each identified signaling pathway is calculated using the right-tailed Fisher’s Exact Test. This significance reflects the probability of the association of molecules from our Patho-DBiT dataset with the canonical pathway reference dataset. Additionally, a graphical summary (Figure 3H and Figure S7F) was generated to provide an overview of the major biological themes in our IPA Core Analysis and illustrate how these concepts interrelate. A machine learning algorithm, relying entirely on prior knowledge, was deployed to score inferred relationships between molecules, functions, and pathways. Networks were constructed from the IPA analysis results using a heuristic graph algorithm.

### Statistical analysis

Statistical analyses were performed using Prism V9 (GraphPad), with the specific tests employed indicated.

## Supplementary Figures and Captions

**Figure S1.**
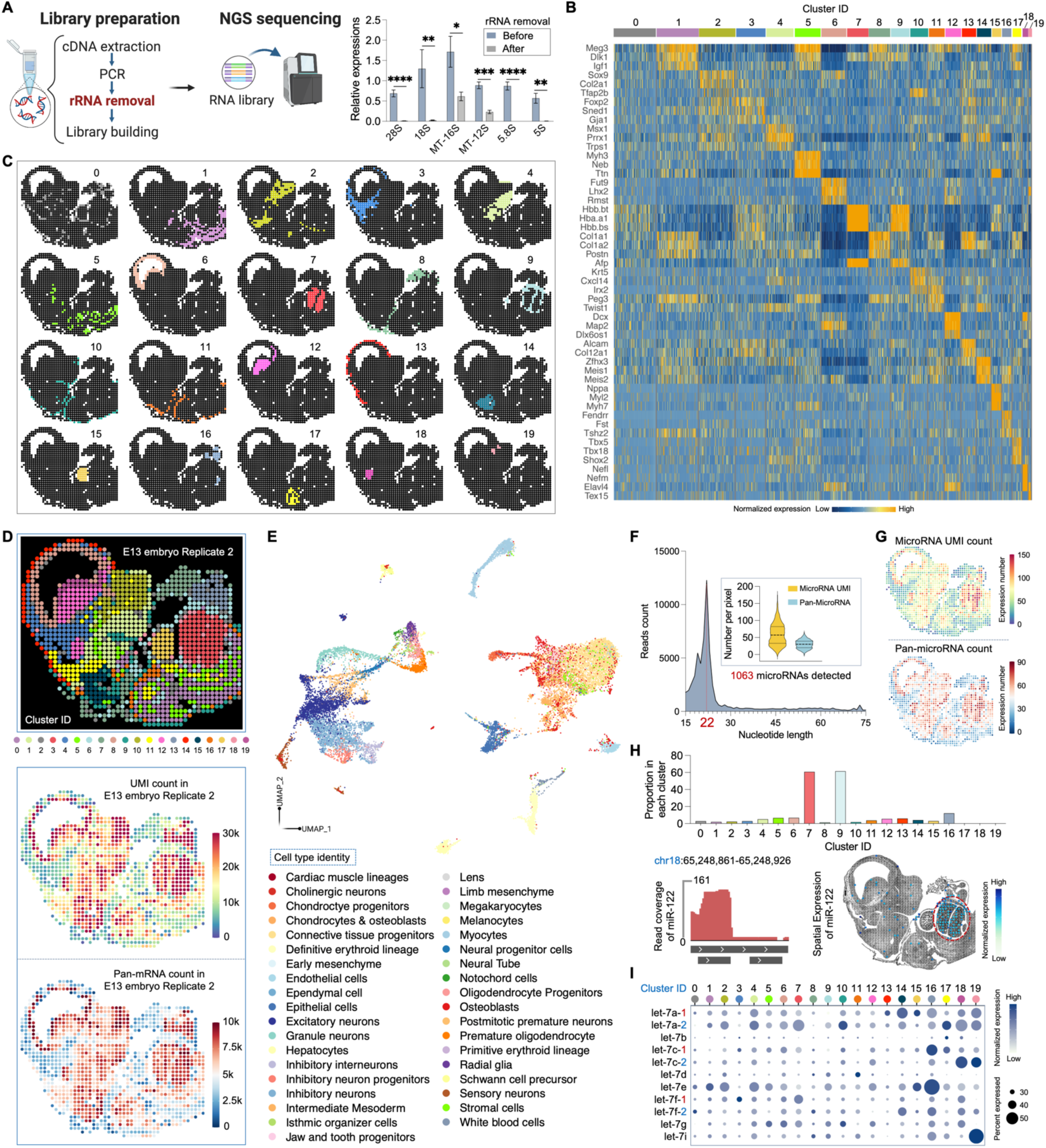
Patho-DBiT performance and spatial microRNA co-profiling in E13 mouse embryo. (A) Left: Patho-DBiT steps post tissue barcoding. Right: qPCR analysis of the rRNA removal efficiency. (B) Heatmap showing top ranked differentially expressed genes (DEGs) defining each cluster. (C) Spatial distribution of identified clusters. (D) Top: unsupervised clustering of the replicate E13 mouse embryo section. Bottom: spatial pan-mRNA and UMI count maps of the replicate section. (E) Cell types identified by integrated clustering analysis of combined scRNA-seq reference dataset (Cao et al., *Nature* 2019) and Patho-DBiT data in E13 mouse embryo. (F) Patho-DBiT demonstrates co-profiling of microRNAs in the E13 mouse embryo. The count of mapped microRNA reads peaks at 22 nucleotides in the dataset. (G) Spatial pan-microRNA and corresponding UMI count maps. (H) Top: expression proportion of the liver-specific miR-122 in each identified cluster. Bottom left: read coverage of miR-122 mapped to the reference genome location. Bottom right: spatial distribution of miR-122. (I) The expression landscape of let-7 family of microRNAs in different spatial clusters. Dot size corresponds to the percentage of pixels expressing the gene, while color shade represents expression level.

**Figure S2.**
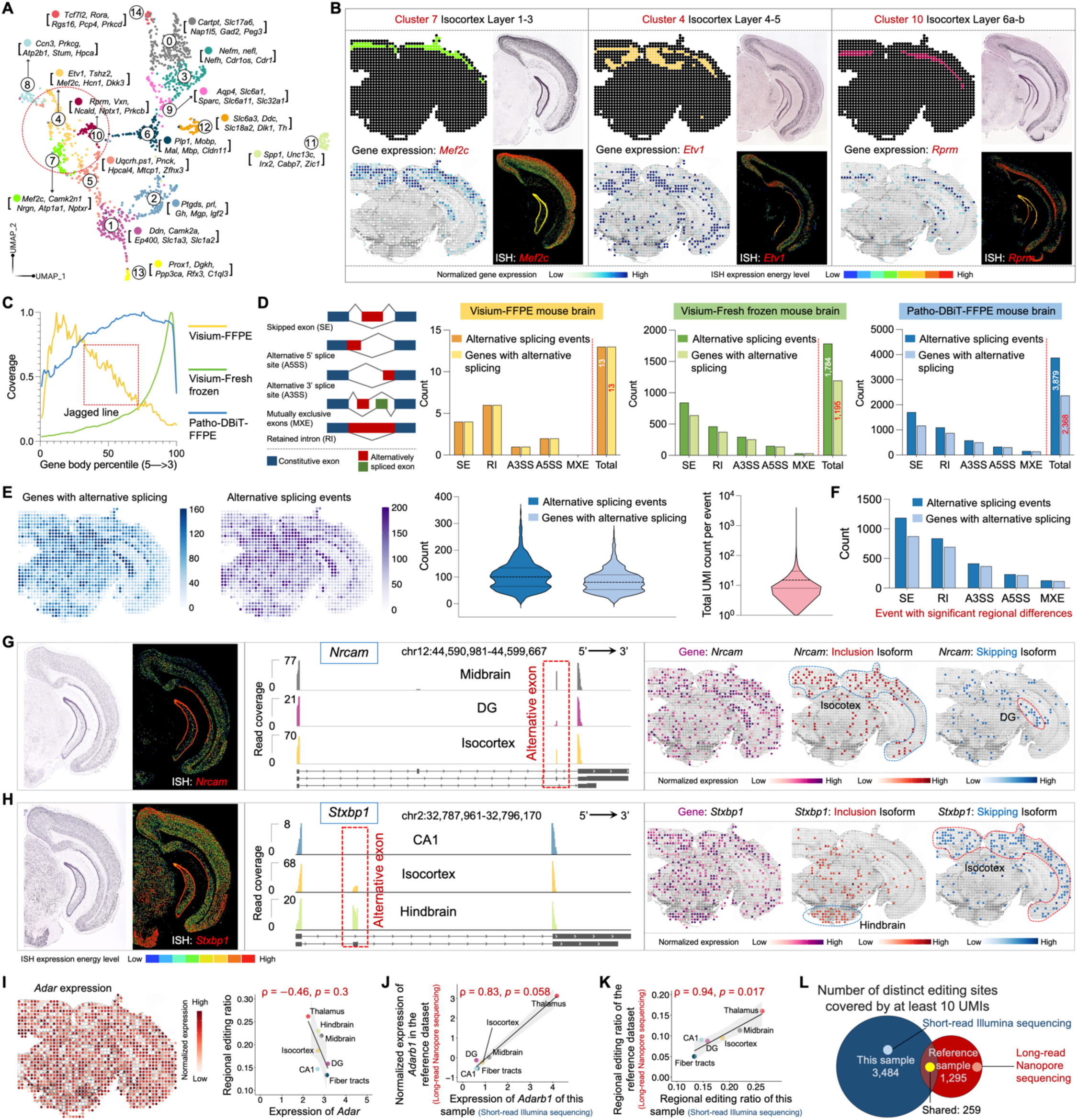
Spatial co-profiling of gene expression, alternative splicing, and A-to-I RNA editing in the mouse brain. (A) UMAP showing the clustering analysis of Patho-DBiT data in FFPE mouse brain section. Top 5 genes defining each cluster are indicated. (B) Spatial distribution of clusters 4, 7, 10 in the isocortex region alongside their principle defining genes. ISH staining and expression images are from the Allen Mouse Brain Atlas. (C) Read coverage along the gene body from 5’ to 3’. Comparison involves Patho-DBiT data and Visium datasets from 10x Genomics on both FFPE and fresh frozen brain sections. (D) Number of detected spliced events and corresponding parental genes within the three datasets listed in (C). (E) Spatial distribution of the detected spliced events and corresponding parental genes of Patho-DBiT data, along with the UMI count distribution per each splicing event. (F) Number of each event type with significant regional differences. (**G** and **H**) Junction read coverage of *Nrcam* (G) and *Stxbp1* (H) splicing event in specific brain regions. Spatial expression patterns of the gene, exon inclusion isoform, and exon skipping isoform are shown. ISH staining and expression images are from the Allen Mouse Brain Atlas. (**I**) Left: spatial *Adar* expression. Right: correlation between the *Adar* expression and the average reginal editing ratio across various brain regions. (**J** and **K**) Correlation between regional *Adarb1* expression (J) or editing ratio (K) detected by short-read Illumina sequencing-based Patho-DBiT and those detected by long-read Nanopore sequencing (Lebrigand et al., *Nucleic Acids Research* 2023). Note, the hindbrain was not included in the brain sample profiled in the reference dataset. (**L**) Venn plot showing the count of distinct editing sites covered by a minimum of 10 UMIs in both datasets.

**Figure S3.**
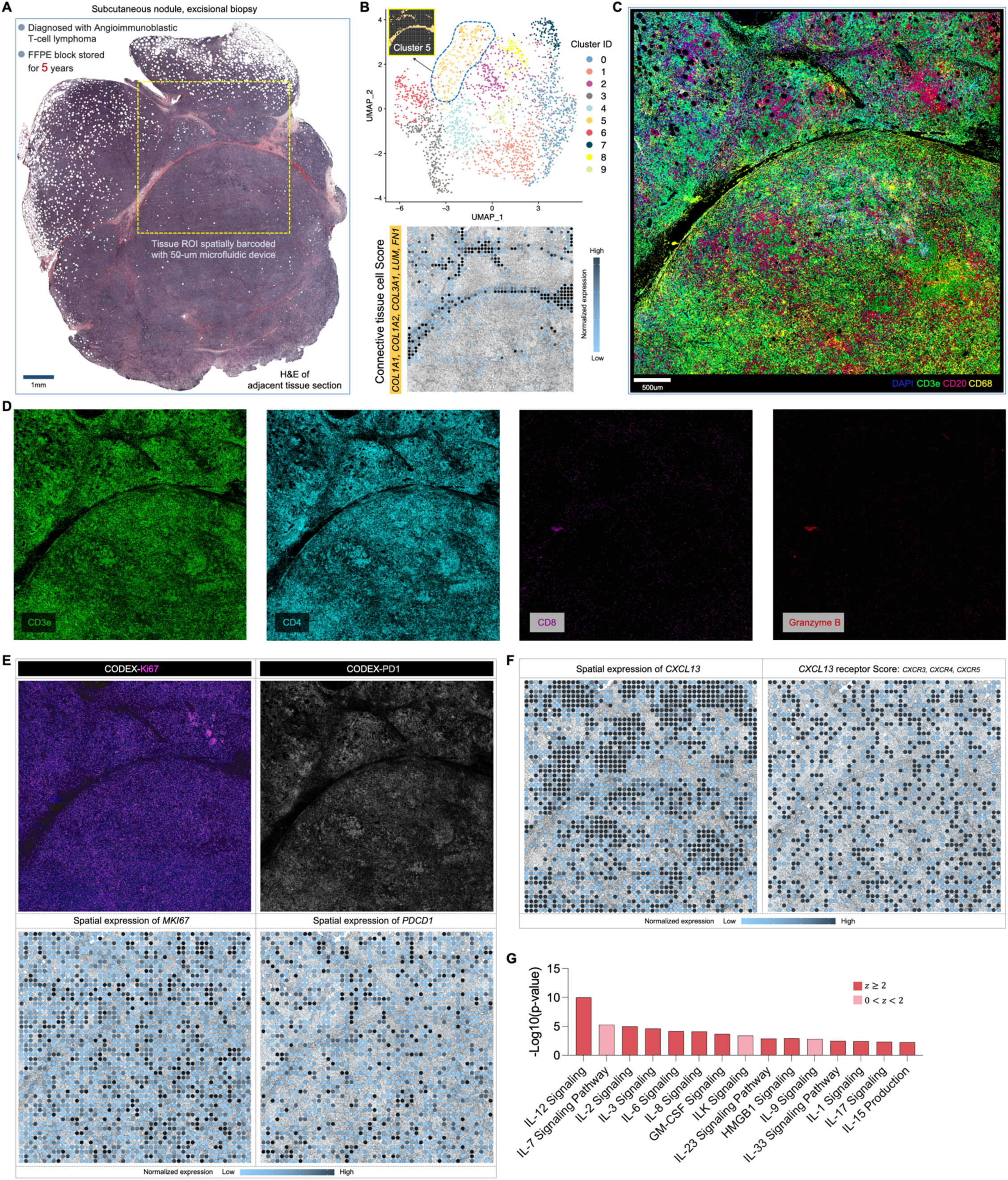
Detection accuracy of Patho-DBiT in the AITL section cross-validated with CODEX. (A) Full scan of the H&E staining of an adjacent section. Yellow square indicates the region of interest (ROI) spatially barcoded with the 50 μm-microfluidic device. (B) Top: UMAP showing the clustering analysis of Patho-DBiT data in the AITL section. The spatial distribution of cluster 5 is indicated. Bottom: spatial expression of the Connective tissue cell Score. Genes defining this module score are listed. (C) Enlarged CODEX image corresponding to the ROI barcoded by Patho-DBiT. (D) CODEX images showing expression of CD3e, CD4, CD8, and Granzyme B. (E) Spatial expressions of *MKI67* and *PDCD1* and the corresponding CODEX images of Ki67 and PD-1. (F) Spatial expressions of *CXCL13* and the *CXCL13* receptor Score. Genes defining this module score are listed. (G) Signaling pathways related to T cell functions in Cluster 0. *z* score is computed and used to reflect the predicted activation level (*z*>0, activated; *z*<0, inhibited; *z*≥2 or *z*≤−2 can be considered significant).

**Figure S4.**
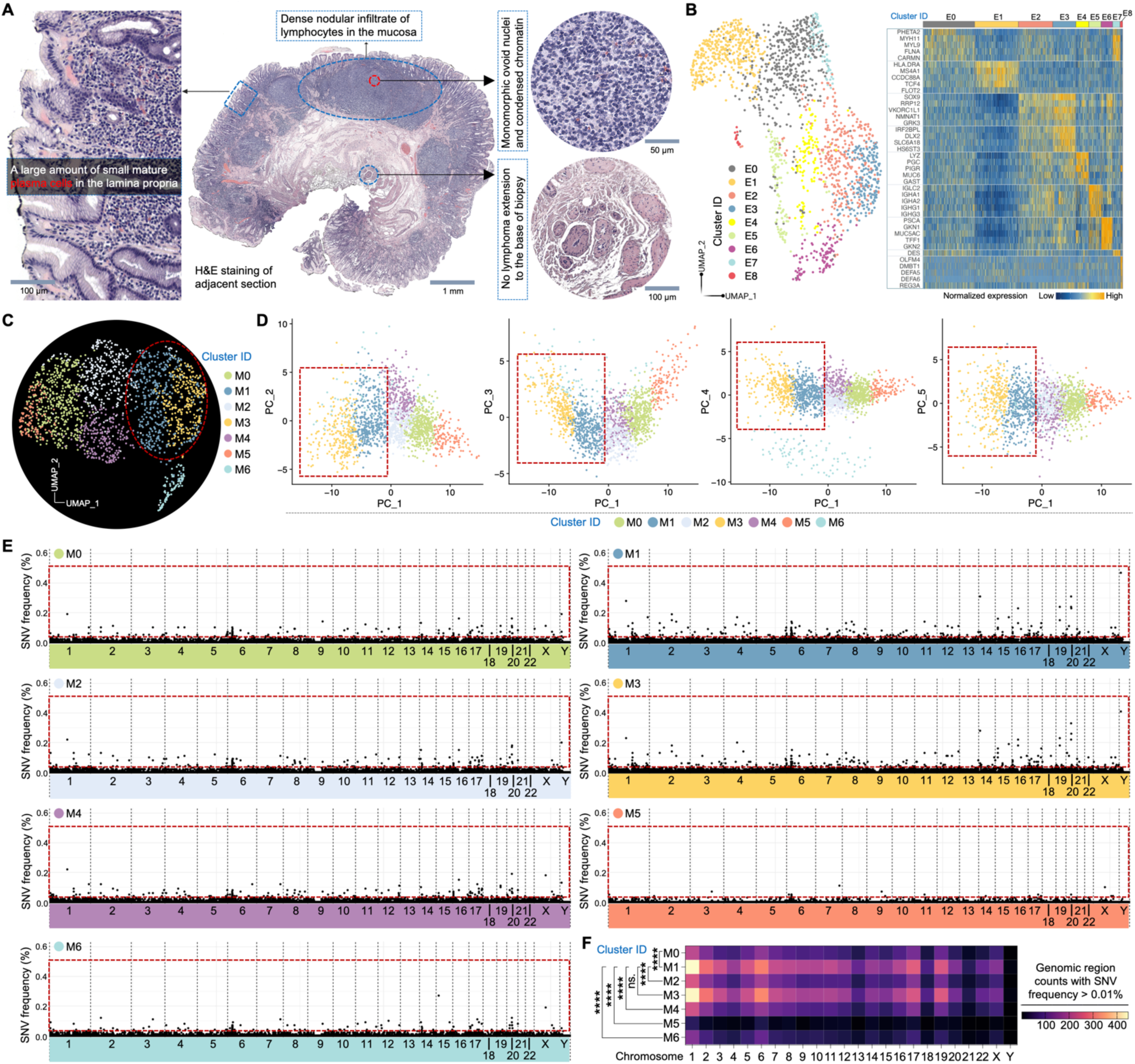
Spatial transcriptome and variation co-profiling of the MALT section. (A) H&E staining of an adjacent section with key histological information annotated by a pathologist. (B) Left: UMAP showing the clustering analysis of Patho-DBiT data in the MALT section. Right: heatmap showing top ranked DEGs defining each cluster. (C) UMAP showing the clustering analysis of spatial SNV matrix. (D) Principal component analysis (PCA) of the identified variation clusters in (B). Top5 PCA components were analyzed. (E) Genome-wide distribution of SNV frequency in each variation cluster. The SNV frequency was counted within sliding genomic regions of 10,000 bp. (F) Count comparison of genomic regions with SNV frequency >0.01% between variation clusters.

**Figure S5.**
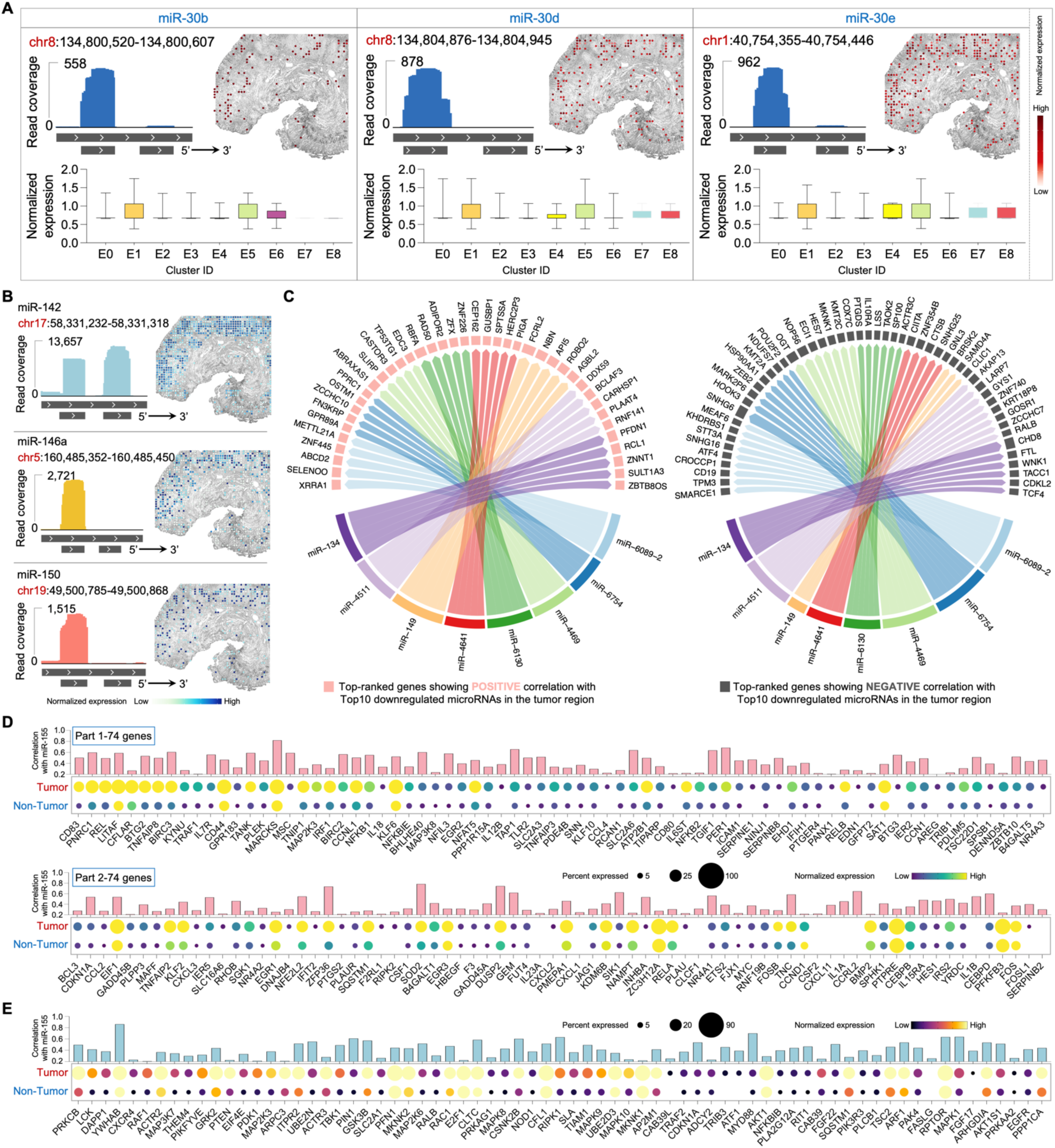
Spatial microRNA profile of the MALT section. (A) Spatial mapping of microRNAs in mature B-cell differentiation regulation. The read coverage mapped to the reference genome location, expression proportion in each identified cluster, and spatial distribution are shown. (B) Spatial mapping of microRNAs enriched in marginal zone B cells or lymphoma. The read coverage mapped to the reference genome location and spatial distribution are shown. (C) Regulatory network between the top 10 downregulated microRNAs and the gene expression in the tumor region. Genes with the highest rankings, demonstrating positive or negative correlations with the microRNAs, were separately illustrated. Edge thickness is proportional to correlation weights. (**D** and **E**) Correlation analysis between miR-155 and each gene in the GSEA-defined NF-κB signaling (D) or PI3K-AKT signaling (E). Enhanced expression of genes participating in both signaling pathways is observed in the tumor region compared to the non-tumor region. The dot size indicates the percentage of pixels expressing the gene, and the color shade represents normalized expression level.

**Figure S6.**
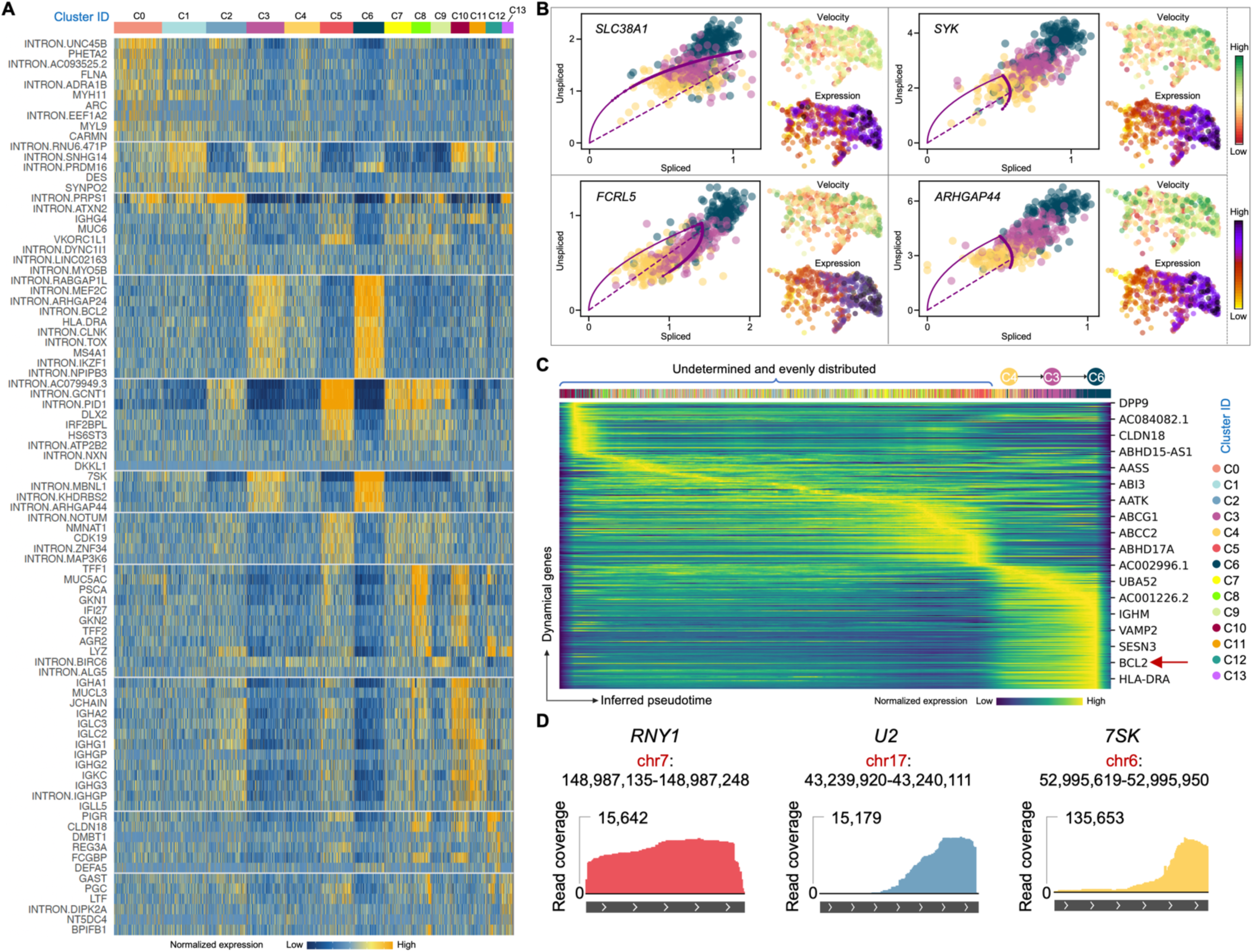
Spatial RNA splicing and pseudotime analysis of the MALT section. (A) Heatmap showing top ranked DEGs defining each cluster identified by clustering analysis of the combined exonic and intronic expression matrix. (B) Phase portraits showing the ratio of unspliced and spliced RNA for more top-ranked genes driving the dynamic flow from cluster C4 to C6, along with their expression and velocity level within the three tumor clusters. (C) Gene expression dynamics resolved along the pseudotime direction showing a clear cascade of transcription of the top-ranked genes. (D) Read coverage mapped to the reference genome location for the selected small RNAs.

**Figure S7.**
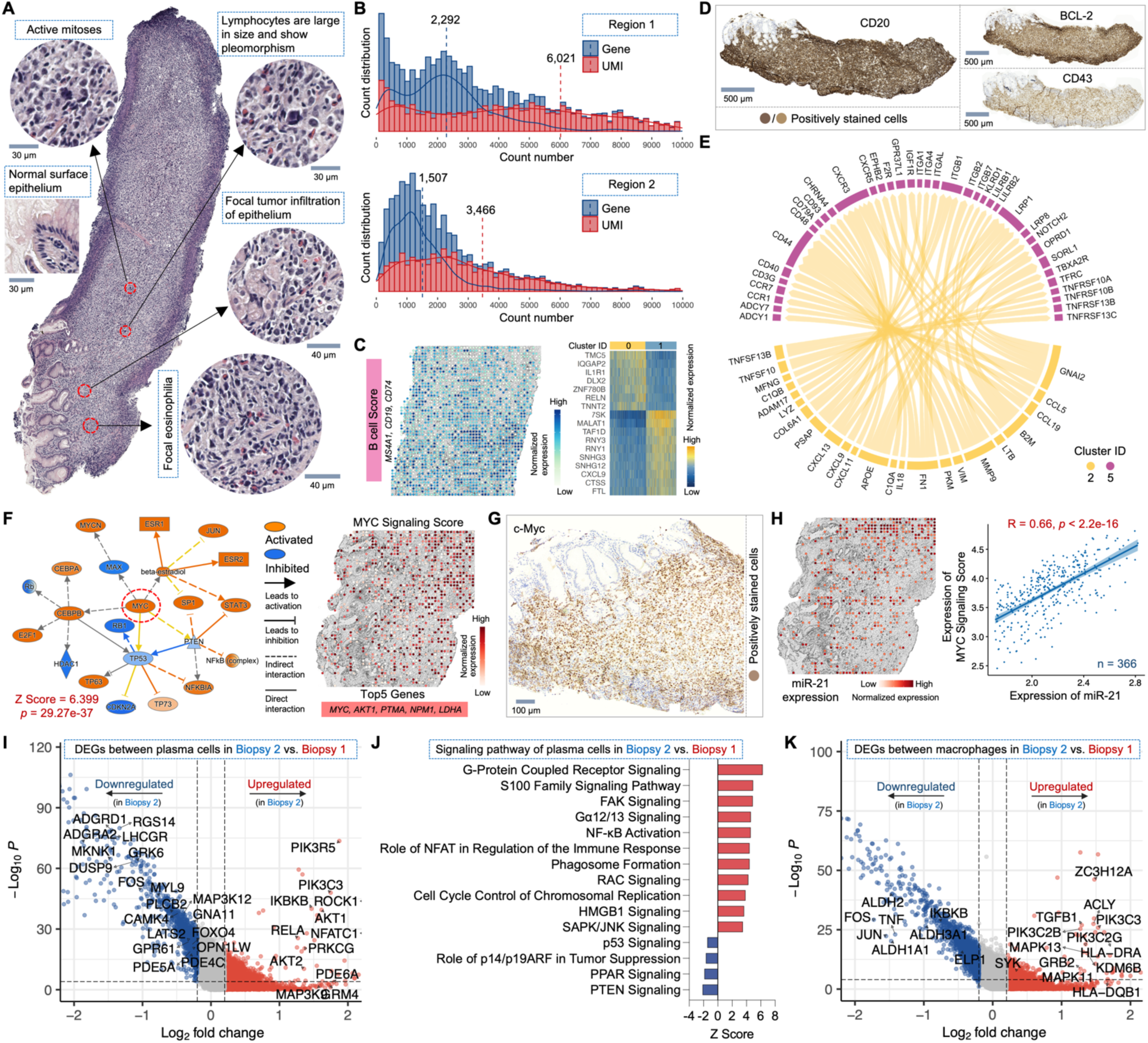
Spatial transcriptome and microRNA analysis of the DLBCL section. (A) H&E staining of an adjacent section with key histological information annotated by a pathologist. (B) Distribution of detected gene/UMI counts per spatial pixel within two DLBCL regions. Only reads mapped to the exonic region were considered. Dashed lines represent the average gene or UMI count level. (C) Left: Spatial expression of the B cell Score in the Region 1 section. Genes defining this module score are listed. Right: heatmap showing top ranked DEGs defining each cluster identified in the Region 1 section. (D) IHC staining of CD20 and canonical markers commonly detected in DLBCL tumor cells (BCL-2 and CD43) on adjacent sections. (E) Ligand-receptor interactions between tumor B cell clusters 2 and 5. Edge thickness is proportional to correlation weights. (F) Left: mechanistic network analysis identifying a significant activation of the master regulator MYC in tumor B cells. Right: spatial expression of the MYC Signaling Score. Top 5 genes defining this module score are listed. (G) IHC staining for c-Myc on an adjacent section. (H) Left: spatial expression map of miR-21. Right: correlation analysis between miR-21 expression and the MYC Signaling Score. The Pearson correlation was calculated across 366 spatial pixels within the tumor region. (I) Volcano plot showing DEGs between plasma cells in DLBCL vs. MALT biopsy. (J) Signaling pathways regulated by DEGs identified in (I). (K) Volcano plot showing DEGs between macrophages in DLBCL vs. MALT biopsy. In (F), and (J), *z* score is computed and used to reflect the predicted activation level (*z*>0, activated; *z*<0, inhibited; *z*≥2 or *z*≤−2 can be considered significant).

